# LTK and ALK regulate neuronal polarity and cortical migration by modulating IGF1R activity

**DOI:** 10.1101/2023.01.29.526107

**Authors:** Tania Christova, Stephanie Ho, Ying Liu, Mandeep Gill, Liliana Attisano

## Abstract

The establishment of axon-dendrite polarity is fundamental for radial migration of neurons, cortical patterning and formation of neuronal circuitry. Here, we demonstrate that the receptor tyrosine kinases, Ltk and Alk, are required for proper neuronal polarization. In isolated primary mouse embryonic neurons, loss of Ltk and/or Alk yields a striking multiple axon phenotype. In mouse embryos and newborn pups, the absence of Ltk and Alk results in a delay in neuronal migration and subsequent cortical patterning. In adult cortices, neurons with aberrant neuronal projections are evident and there is a disruption of the axon tracts in the corpus callosum. Mechanistically, we show that loss of Alk and Ltk increases cell surface expression and activity of the insulin-like growth factor 1 receptor (Igf-1r), which acts to activate downstream PI3 kinase signalling to drive the excess axon phenotype. Thus, our data reveal Ltk and Alk as new regulators of neuronal polarity and migration whose disruption results in behavioural abnormalities.

## Introduction

Neuronal polarization, namely the specification of axon and dendrites, is critical for the development and proper functioning of the brain (Arimura & Kaibuchi, 2007, Barnes & Polleux, 2009, Takano, Xu et al., 2015). During early development of the CNS, neural progenitors located in the ventricular zone transition from a multipolar to bipolar state, which enables subsequent radial migration and formation of defined layers that make up the cortex (Noctor, Flint et al., 2001). In adults, the proper neuronal morphology is critical for signal transmission and for formation of neuronal circuitry that drives memory, behavior and all key brain activities (Arimura & Kaibuchi, 2007, Takano et al., 2015). In line with this, alterations in neurogenesis, neuronal migration or morphology can result in neurodevelopmental defects that lead to a variety of neurological and neuropsychiatric disorders (Romero, Bahi-Buisson et al., 2018). A wide variety of signalling pathways and factors have been identified as regulators of neuronal polarization including Rho GTPases, phosphatidylinositol 3-kinase (PI3K), MAPK pathway components, Wnts and Receptor Tyrosine Kinases (RTKs) (Arimura & Kaibuchi, 2007, Lewis, Courchet et al., 2013, Takano et al., 2015).

Leukocyte tyrosine kinase (LTK) and Anaplastic lymphoma kinase (ALK) are closely related Receptor Tyrosine Kinases (RTKs) belonging to the insulin receptor superfamily (Palmer, Vernersson et al., 2009, Roskoski, 2013, Weiss, Xue et al., 2012). ALK was initially described as oncogenic fusion protein in anaplastic large cell lymphomas (ALCL) (Morris, Kirstein et al., 1995, Shiota, Fujimoto et al., 1994) and since then, aberrantly activated ALK arising from protein fusions, point mutations or overexpression have been described in many cancers including neuroblastoma (Hallberg & Palmer, 2016, Janoueix-Lerosey, Lopez-Delisle et al., 2018, Palmer et al., 2009). In general, inappropriate engagement of characteristic RTK downstream effectors are thought to mediate the pro-oncogenic activities of these fusion and/or mutant forms of ALK (Hallberg & Palmer, 2016, Palmer et al., 2009). Although less studied, LTK dysregulation also has consequences for cancer progression (Izumi, Matsumoto et al., 2021, Muller-Tidow, Schwable et al., 2004). Accordingly, effective small molecule inhibitors of ALK/LTK have been developed for clinical use, primarily for non-small cell lung cancers (Awad & Shaw, 2014, Chuang & Neal, 2015). Metastasis to the brain is frequently observed in patients with ALK mutations, an observation that prompted the development of second round therapeutics, such as lorlatinib, that have successfully addressed the inadequate central nervous system penetration of the first-round inhibitor, crizotinib (Awad & Shaw, 2014, Chuang & Neal, 2015). Ongoing clinical trials have confirmed intracranial activity, however, some changes in cognitive function, mood and speech in lorlatinib-treated patients have been reported (Solomon, Besse et al., 2018).

Outside of oncogenic contexts, both ALK and LTK are predominantly expressed in the central and peripheral nervous systems (Iwahara, Fujimoto et al., 1997, Janoueix-Lerosey et al., 2018, Weiss et al., 2012). However, in marked contrast to the extensive literature on ALKs in cancer, the normal role of the receptors is much less understood. ALK and/or LTK have been shown to play a role in the development and function of the nervous system. In Zebrafish, LTK promotes the survival and migration of neural crest cells and is required for the specification of the neural crest-derived pigment cells, called iridophores (Fadeev, Mendoza-Garcia et al., 2018, Lopes, Yang et al., 2008) while ALK is required for proper neuronal differentiation and survival in the CNS (Yao, Cheng et al., 2013). In Drosophila, ALK is required for neuronal circuit assembly in the developing retina and neuromuscular junction, in sparing of nervous system growth during nutrient deprivation in larva (Cheng, Bailey et al., 2011), and in adult learning (Gouzi, Moressis et al., 2011). In mice, overexpression of an activated Alk receptor during embryogenesis results in a disruption of the differentiation of neural crest progenitors (Vivancos Stalin, Gualandi et al., 2019) and in an Alzheimer’s model of tau proteinopathy, that aberrant activation of Alk leads to the abnormal accumulation and aggregation of phosphorylated Tau and modulates mouse behaviour (Park, Choi et al., 2021). Mice deficient in Alk and/or Ltk are viable and lack gross morphological alterations, but do display several subtle behavioural abnormalities, responses to ethanol (Bilsland, Wheeldon et al., 2008, Lasek, Lim et al., 2011, Weiss et al., 2012), hypogonadotropic hypogonadism in male mice (Witek, El Wakil et al., 2015) and resistance to diet-induced obesity (Orthofer, Valsesia et al., 2020).

The ligands for LTK and ALK, named ALKAL1 and ALKAL2 (previously referred to as FAM150A/B or Augmentor β/α), were identified relatively recently (Guan, Umapathy et al., 2015, Reshetnyak, Murray et al., 2015, Zhang, Pao et al., 2014), and a series of recent studies have provided important structural insights into how the ligands engage and activate the receptors (De Munck, Provost et al., 2021, Li, Stayrook et al., 2021, Reshetnyak, Rossi et al., 2021). The ligands have also been shown to be important in CNS development and cancer, acting to specify iridophores in Zebrafish (Mo, Cheng et al., 2017), functioning in the hypothalamus to control body weight in mice (Ahmed, Kaur et al., 2022) and in driving tumorigenesis in MYCN-driven non-mutant ALK neuroblastomas (Borenas, Umapathy et al., 2021).

In this study, we sought to understand the physiological role of LTK and ALK in the central nervous system, specifically during cortical development using mouse models. We observed that in the absence of Ltk and Alk, neural progenitor populations, neuronal migration and subsequent patterning of the cortical layers is disrupted in developing mice and that isolated primary embryonic cortical neurons displayed a multiple axon phenotype. We next investigated the underlying mechanism for the aberrant neuronal phenotype and demonstrate that loss of Alk and Ltk increases cell surface expression and activity of insulin-like growth factor 1 receptors (Igf-1r) that in turn activate downstream PI3 kinase signalling to drive the multiple axon phenotype. Taken together, our data reveal LTK/ALK as new regulators of neuronal morphology and migration. This study, thus suggests that it will be important to consider the potential impact on CNS and brain functions in cancer patients being treated with pharmacological inhibitors of ALK/LTK activity.

## Results

### Ltk and Alk are expressed in the embryonic and adult cortex

ALK and LTK have been extensively studied in cancer, but ittle is known of the normal biological role of these receptors. Thus, to gain insights into the normal function of Ltk and Alk, we first examined the localized expression of these receptors using single molecule *in situ* mRNA analysis in the developing cortex. At E15.5, Ltk showed broad expression in the superficial layers (ie cortical plate: CP and subcortical plate: SCP) and in intermediate and ventricular/subventricular zones (VZ/SVZ; Fig. S1A and B). Alk expression was also broadly localized, but the highest expression was observed in the proliferating VZ that was marked by Sox2 (Fig. S1A and B). Both receptors were also expressed in several regions outside the neocortex (Fig. S1C-F) consistent with previous findings (Janoueix-Lerosey et al., 2018, Weiss et al., 2012). Analysis of expression by real-time PCR revealed that both receptors were expressed at E15.5 in the cortex and brain but in differing temporal patterns, with Alk expression maximal at E15.5 and then declining to adulthood, whereas Ltk showed a dramatic enhancement in adults, particularly within the cortex (Fig. S1G).

ALKAL1 and ALKAL2 (also known as FAM150A/Augmentor β and FAM150B/Augmentor α, respectively) are the ligands for LTK and ALK (Guan et al., 2015, Reshetnyak et al., 2015, Zhang et al., 2014). Single-molecule *in situ* hybridization confirmed that Alkal2 was widely expressed in the E15.5 cortex in regions overlapping the Ltk and Alk expression domains including the cortical plate (CP) and ventricular zone (Fig. S1A-F). Analysis by real-time PCR revealed that Alkal2 was expressed in the mouse embryonic, post-natal and adult cortex and brain at roughly equivalent levels (Fig. S1G). In contrast, Alkal1 expression was not detected by qPCR or *in situ* in the mouse brain (not shown), consistent with previous reports (Ahmed et al., 2022) and data in the Protein Atlas database (Uhlen, Fagerberg et al., 2015, www.proteinatlas.org) Altogether these data show that Alk and Ltk and their ligand, Alkal2 are expressed throughout the embryonic and adult cortex.

### Loss of Ltk and Alk disrupts neuronal cell migration and cortical layer patterning in the developing cerebral cortex

To investigate the role of Ltk and Alk in normal physiology, we generated double knockout (*DKO*) mice constitutively lacking both Ltk and Alk (*Ltk^-/-^Alk^-/-^*) by crossing homozygous *Ltk^-/-^* and *Alk^-/-^* single knockout animals. *Ltk* and *Alk* single and *DKO* mice were viable and fertile as previously reported (Bilsland et al., 2008, Weiss et al., 2012). To investigate the consequences of the loss of Ltk/Alk in the cortex, we first examined the status of progenitor cells in the neurogenic niche. During early cortical development, neural stem cells (NSCs) proliferate symmetrically in the VZ while at the onset of neurogenesis, apical radial glial progenitor (aRG) cells in the VZ divide asymmetrically to produce intermediate neuronal progenitor (INP) cells and post-mitotic neurons, both of which lack Sox2 expression (Gotz & Huttner, 2005). Thus, we examined the effect of the loss of *Ltk* and/or *Alk* on progenitor cell populations in E15.5 mice. WT mice displayed the characteristic pattern of Pax6/Sox2 positive aRG cells in the VZ and Tbr2+ INP in the SVZ (Fig.1A and Fig. S2A). However, in single and DKO mice, while the total number of cells (DAPI+) and number of Pax6+ cells was similar to WT, a marked reduction in both Sox2+ cells, which mark multipotent progenitors and Tbr2+ cells, which indicate INPs, was observed with DKO mice displaying the most dramatic reduction (Fig. 1A, Fig S2A). Thus, loss of Ltk/Alk results in a disruption of neurogenesis.

**Figure 1.**
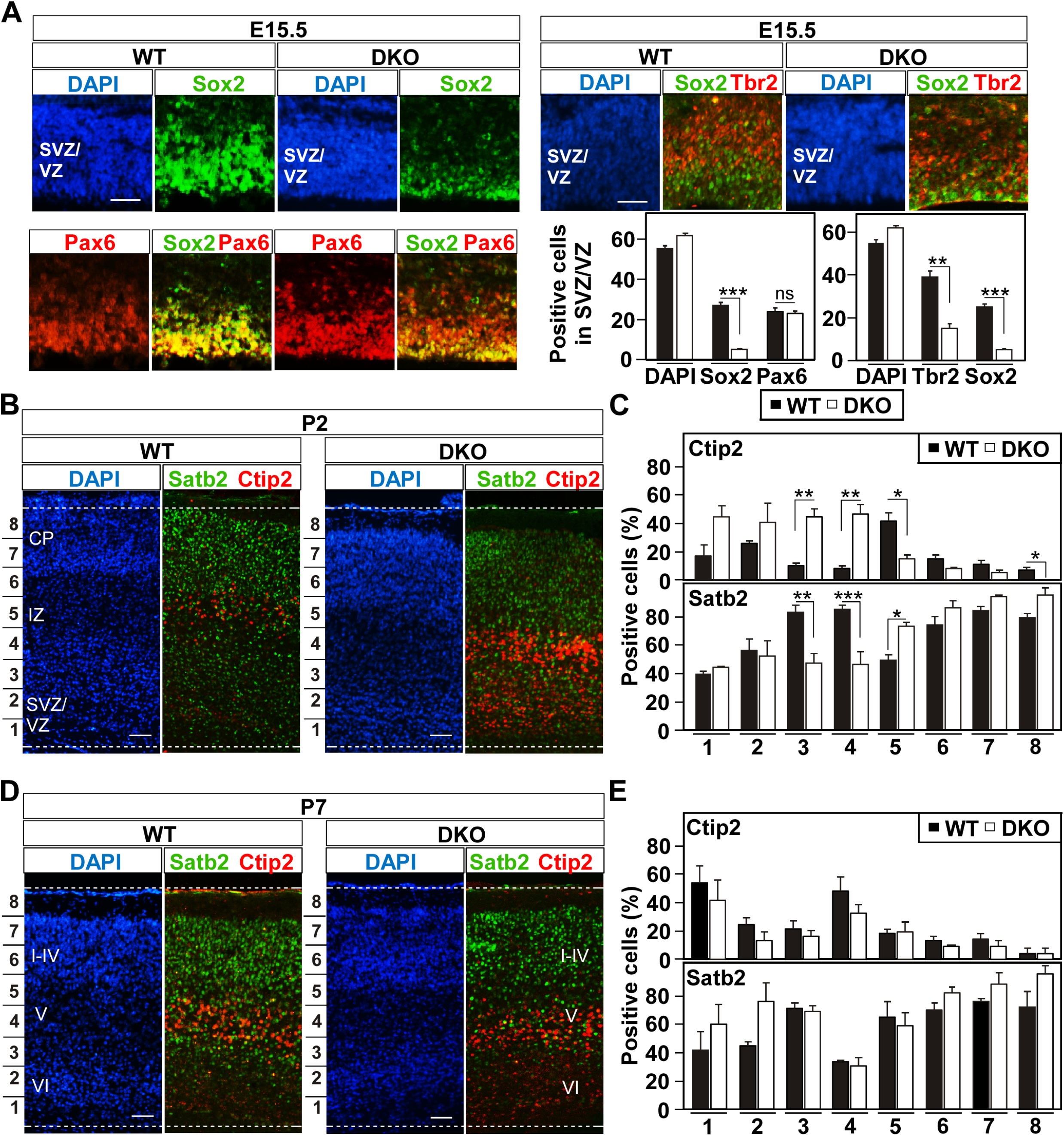
Disruption of Ltk and Alk alters early cortical plate patterning. **(A)** Coronal sections of E15.5 wild-type (WT) and DKO developing cortices were co-stained with either anti-Sox2 (green) and anti-Pax6 (red) or with anti-Sox2 (green) and anti-Tbr2 (red) and both counter-stained with DAPI (blue). Quantitation of total DAPI+ cells and Sox2+, Pax6+ or Tbr2+ progenitor cells in the SVZ/VZ are plotted as the mean +/- SEM of 3 independent experiments. **(B and D)** Coronal sections of WT and DKO developing cortices stained with DAPI (blue) and co-stained with anti-Ctip2 (red) and Satb2 (green) at P2 and P7. **(C and E)** Quantitation of Ctip2+ and Satb2+ progenitor cells as a percent of total DAPI+ cells in each of 8 bins (marked on left) within the cortex (dashed lines) are plotted as the mean +/- SEM of 3 independent experiments. Statistical significance: ***p<0.001, ** p<0.01, *p<0.05, Student’s t-test. The location of the cortical plate (CP), intermediate zone (IZ), subventricular/ventricular zone SVZ/VZ in P2 mice and Layers I-VI in P7 WT mice are indicated. Scale bar, 50 µm.

Newly born neurons migrate through the intermediate zone (IZ) and give rise to five cortical layers distinguished by the expression of distinct transcription factors that control cortical neuronal identities and properties (Molyneaux, Arlotta et al., 2007, Uzquiano, Gladwyn-Ng et al., 2018). Thus, to investigate whether postmitotic regions of the developing cortex were also affected by the loss of Ltk and Alk, we next examined the localization of two layer-specific markers, Ctip2, and Satb2 in WT and knockout mice. At post-natal day 2 (P2), a timepoint in which neurogenesis and migration have primarily subsided, WT Ctip2+ cells are primarily localized to the IZ (bins 5/6) in WT mice, while in DKO mice, the percent of Ctip2+ cells in this region was notably reduced with many Ctip2 positive cells also localized to the lower layers (bins 1-4, marked SVZ/VZ in WT mice: Fig. 1B and C). By P7, when neuronal migration is largely completed, the localization of Ctip2+ neurons in DKOs was more similar to the WT, though some Ctip2 positive cells were still observed in the lowest layers (ie bins 1-4, labelled Layer VI in WT mice; Fig. 1D and E). In the case of Satb2, a reduction in percent of positive cells in some of the lower layers was observed in mutants at P2 (bins 3 and 4), possibly due to the high proportion of Ctip2+ cells in this region, that was resolved by P7 (Fig. 1B-E). The improvement was also evident in Nissl-stained sections of adults where DKO cortices did not display any cytoarchitectural defects (Fig S3). Analysis of single *Ltk^-/-^*and *Alk^-/-^* mice at P2 and P7 revealed similar, though more modest disruptions in the localization of Ctip2 and Satb2 positive neurons (Fig. S2C-F). Given that Ctip2 positive neurons were present but showed altered timing of proper localization, our observations suggest that loss of *Ltk* and *Alk* results in a delay in migration rather than in a disruption in neuronal subtype specification.

We next sought further evidence that the alterations in patterning of the cortical layers in the absence of Ltk/Alk were due to a disruption in the migration of nascent neurons. For this, embryos were injected with BrdU at E14.5 and were then fixed and stained at E17.5, P2 and P7 (Fig. 2). Quantitative analysis revealed that in E17.5 wild-type mice, the majority of BrdU-positive neurons had migrated towards the superficial layers, spanning the entire cortex. In contrast, in single *Ltk^-/-^* or *Alk^-/-^* and in *DKO* mice, there was a marked decrease in BrdU-positive neurons reaching the middle and superficial layers (bins 5 – 8: Fig 2A and B). In P2 WT brains, most of the BrdU-positive neurons were found in bin 6 – 8 (ie layers I-III), whereas in the *DKOs*, most of the BrdU-positive neurons were scattered throughout the cortex. By P7, the majority of BrdU-positive neurons were found in the superficial layers in both DKO and WT brains (Fig 2C and D). This apparent delay in migration recapitulates the delayed patterning observed using layer specific markers, Ctip2 and Satb2 shown in Fig. 1B-E. Taken together these studies demonstrate a role for Alk/Ltk in promoting neuronal migration during patterning of the developing cortex.

**Figure 2.**
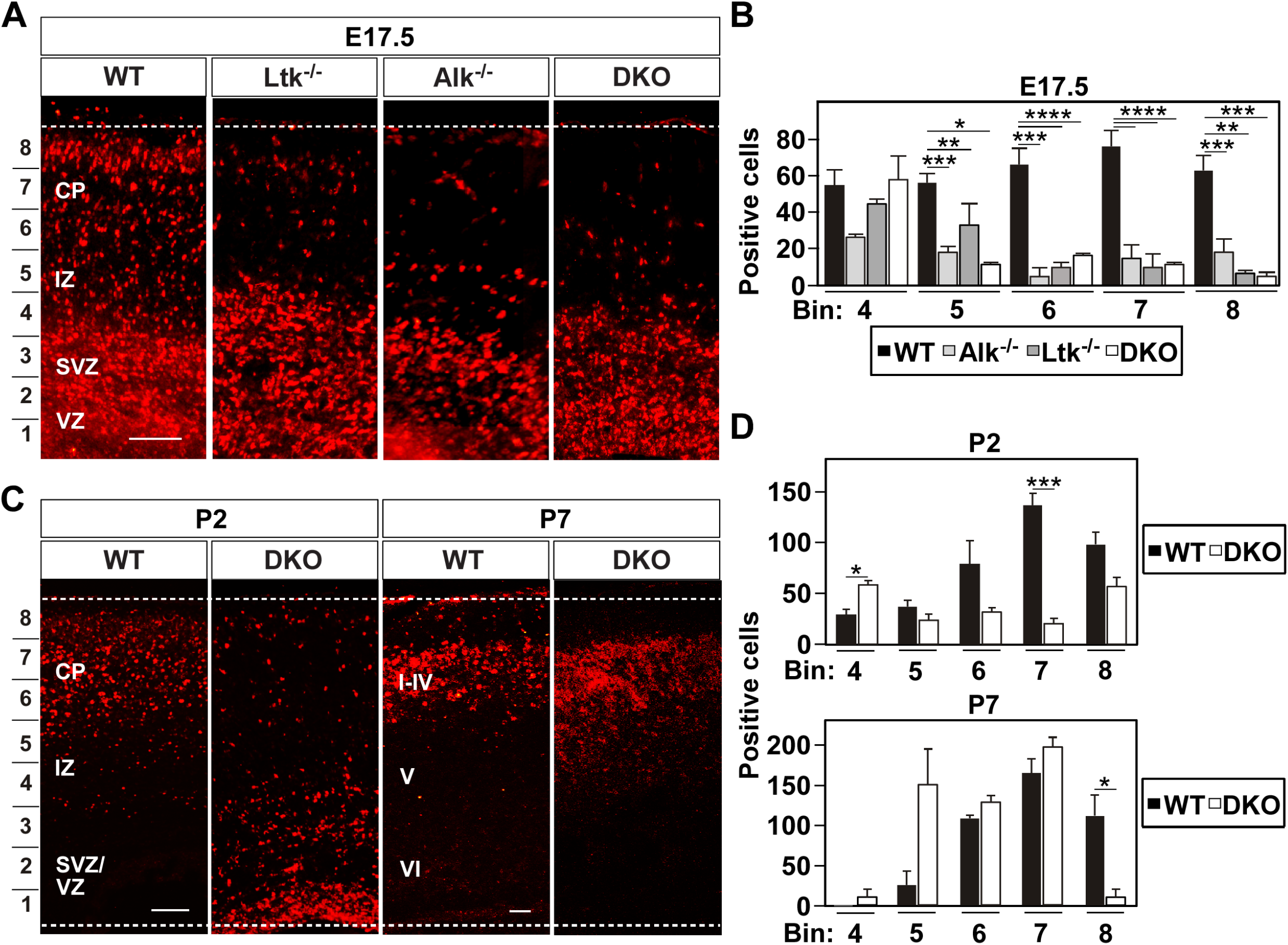
Loss of Ltk/Alk leads to delayed radial migration. Wild type (WT), single (Ltk-/- and Alk-/-) and double (DKO) knockout embryos were injected with BrdU on day E14.5 and fixed cortices analyzed at E17.5, P2 and P7. **(A and C)** Coronal sections of BrdU labeled WT and mutant cortices at the indicated ages. **(Band D)** Quantitation in E17.5 **(B)** or P2 and P7 **(D)** cortices of BrdU-positive cells in the indicated bins (marked on left) is plotted as the mean +/- SEM from 3 (DKO, Ltk-/-) or 2 (WT, Alk-/-) independent experiments **(B)** or 3 DKO and WT independent experiments **(D).** Statistical significance: ****p<0.0001, ***p<0.001, ** p<0.01, *p<0.05 by Student’s t-test. The location of the cortical plate (CP), intermediate zone (IZ), subventricular/ventricular zone SVZ/VZ in E17.5 and P2 mice and Layers I-VI in P7 WT mice are indicated. Scale bars are 50 µm.

### Ltk/Alk regulate neuronal polarity in cortical neurons

Proper neuronal polarization is essential for migration of nascent neurons (Arimura & Kaibuchi, 2007). Thus, to explore a potential role for Ltk/Alk in neuronal polarization, we turned to freshly isolated and plated primary embryonic cortical neurons, a model system that recapitulates the *in vivo* acquisition of neuronal polarity (Fig. 3A) in which a single axon emerges from multiple indistinguishable neurites (Barnes & Polleux, 2009, Bradke & Dotti, 2000). We confirmed by single molecule RNA *in situ* that the majority (70%) of isolated neurons expressed both Ltk and Alk, roughly 20% express only one receptor and about 10% express neither (Fig. S4A). We observed that primary cortical neurons isolated from single *Ltk^-/-^* or *Alk^-/-^* mice displayed a marked increase in the number of axons, visualized by staining with axonal marker Tau-1 that were negative for the dendritic marker, MAP2, indicating axonal identity (Fig. 3B). Simultaneous loss of both Ltk and Alk (ie *DKO* neurons) did not further enhance the multiple axon phenotype, suggesting that these receptors do not act redundantly in this context, but rather may both be required for proper neuronal polarization function. Conversely, overexpression of HA-tagged LTK (LTK-WT-HA) inhibited axon formation in wild-type neurons, while in *DKO* neurons, ectopic expression of LTK not only reversed the aberrant multiple axon phenotype, but also restored the wild-type single axon morphology (Fig. 3C and S4B). We confirmed these observations by testing the effect of abrogating the expression of *Ltk*, *Alk* or both in wild-type neurons using siRNAs. Similar to neurons isolated from knockout mice, we observed a marked increase in the number of axons upon loss of Ltk, Alk or both (Fig. S4C and D) indicating that the effect observed in neurons from KO mice was not due to the emergence of indirect, long term compensatory mechanisms. As an alternative to genetic approaches, we also treated wild-type embryonic cortical neurons with two small molecules, TAE684, a potent and highly specific inhibitor of ALK (Galetta, Rossi et al., 2012) and crizotinib, a drug in clinical trials for the treatment of ALK-fusion protein positive cancers (Rodig & Shapiro, 2010), which also inhibits LTK (Centonze, Reiterer et al., 2019). Neurons treated with either TAE684 or crizotinib also displayed the multiple axon phenotype (Fig. 3D). Taken together, these results show that both LTK and ALK are important for proper neuronal polarization in murine cortical neurons and that either abrogation of expression or chemical inhibition of their activities induces the formation of multiple axons.

**Figure 3.**
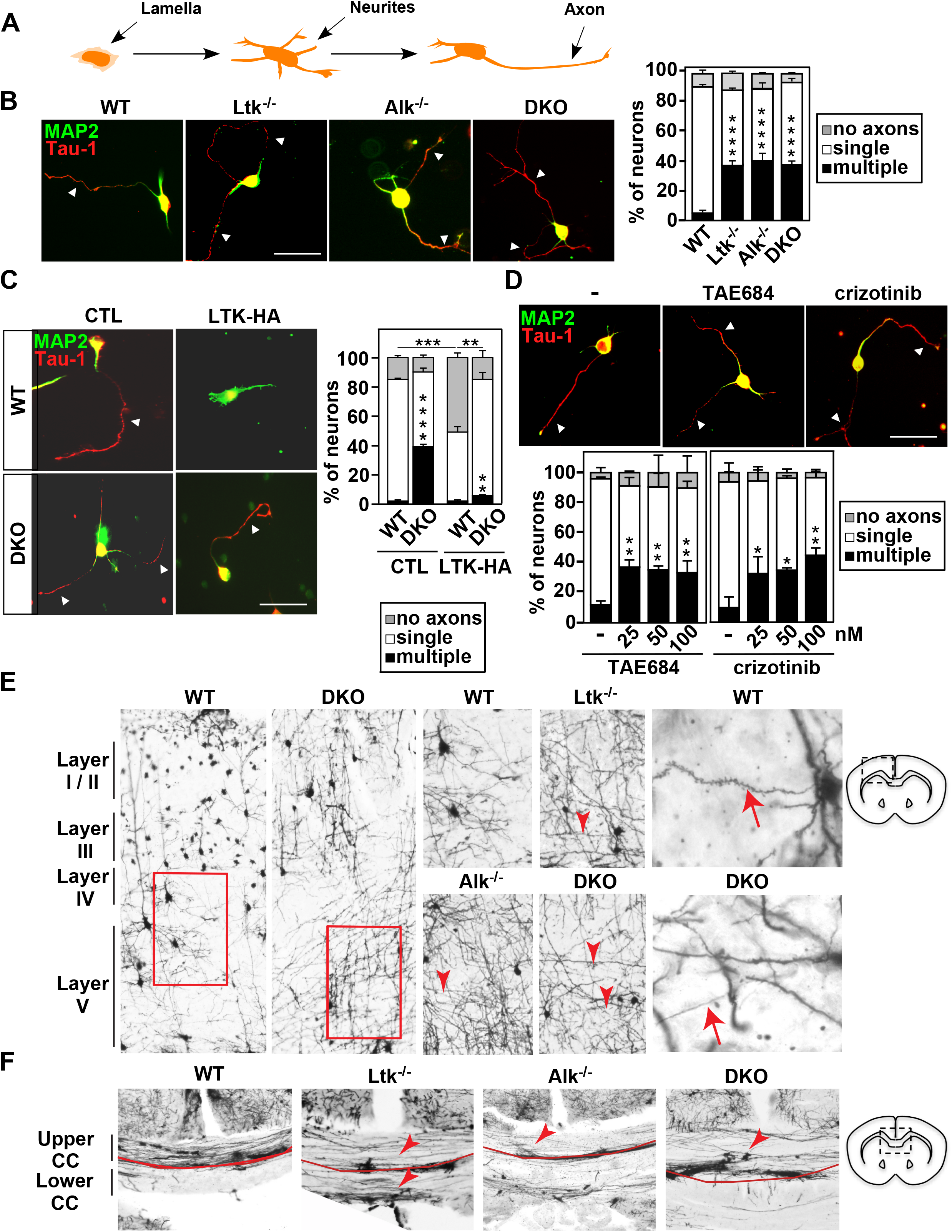
Alterations in Ltk and Alk expression or activity disrupts neuronal polarity in cortical neurons and results in defects in neuronal morphology in the brains of knockout mice. (**A**) A schematic illustrating the development of embryonic cortical neurons cultured in vitro. (**B**) Dissociated E16 cortical neurons were cultured for 38 h and then fixed. Representative images of cortical neurons from WT, single or DKO mice, stained for Tau-1, an axonal marker (red) and MAP2, a dendritic marker (green). (**C**) Representative images of dissociated WT and DKO cortical neurons, electroporated with plasmids encoding EGFP alone or together with HA-tagged LTK-WT. Cells were fixed 38 h after plating and then stained for Tau-1 (axons, red) and MAP2 (dendrites, green). Quantitation of the percent of neurons with multiple, single and no axons in EGFP positive neurons is plotted as the mean ± SEM (n=150 neurons) from 3 independent experiments. (**D**) Representative images of WT cortical neurons, treated with the ALK inhibitors, TAE684 and crizotinib or DMSO as a control at 4 h after plating. Neurons were stained for Tau-1 (red) and MAP2 (green). (**B, D**) Quantitation of the percent of neurons with multiple, single and no axons is plotted as the mean +/- SEM of 150 of neurons per condition from 3 independent experiments. Statistical significance: ****p<0.0001, ***p<0.001, **p<0.01, *p<0.05 by one–way ANOVA, Dunnett’s test. (**E**) Golgi stains of adult cortices in wild-type, single (Ltk^-/-^ and Alk^-/-^) and double (DKO) knockout mice show severe defects in neuronal projections and the callosal tract. Representative coronal sections of cortical layers I-V in WT and DKO mice (left), Layers IV-V in WT, single and double knockout cortices (center) and higher magnification images of WT and DKO neurites (right). Examples of disrupted directional growth and extension of projections in pyramidal neurons in knockout mice are marked (red arrowheads). (**F**) A representative coronal sections of the medial region of corpus callosum in wild-type and mutant cortices. Red line delineates the upper and lower regions of the corpus callosum (CC) with higher axon density within the upper corpus evident. Prominent defects in the callosal fibers including substantial deviations and incoherent projections from the major tract in knockout mice compared to WT mice are marked (red arrowheads). Schematics of the regions represented in images are shown on the right. CC: Corpus callosum, Py: pyramidal neurons.

### *Ltk* and *Alk* knockout mice display defects in neuronal morphology and axon tracts

To investigate whether mice lacking Ltk/Alk display any morphological changes *in vivo*, we visualized neurons in the adult brain using Golgi staining, an approach that enables visualization of individual neurons. The gross overall morphology of wild-type, single or double knockout cortices was similar (Fig. 3E and S3B). In wild-type cortices, analysis of layer IV/V revealed that individual pyramidal neurons were fully developed with elaborated dendritic trees, including dendrites with numerous spines and an apparent single axon (distinguished by an absence of spines) extending to the superficial layers of the mature cortex (Fig. 3E and S3B). In contrast, pyramidal neurons in single and double knockout mice exhibited an abnormal pattern of neurites that included aberrant horizontal projections across the cortical layers (red arrows) that were more prominent in *DKO*s. Of note, within the corpus callosum (CC), abnormalities in the axon tracts were observed in both single and double knockout mice (Fig. 3F). Thus, although adult mice display an overall normal cortical layering morphology, loss of either Ltk alone, Alk alone or both, disrupts the morphology of individual neurons and the proper establishment of axon tracts.

### Increased anxiety and impaired problem-solving behaviour in *DKO* mice

Defects in axons and axon tracts caused by Ltk/Alk deficiency may impact neuronal connectivity and function. Thus, we undertook a series of tests to explore whether these mice display any behavioral abnormalities related to neurodevelopmental disorders, such as deficits in cognitive function and anxiety. We first performed the Light-Dark test, which is based on the innate aversion of rodents to brightly illuminated areas and on their spontaneous exploratory behavior in response to mild stressors (Bourin & Hascoet, 2003). Mice were placed in an open chamber with bright light and the time to discover the adjacent enclosed dark chamber as well as the frequency of exit and re-entry into the dark was determined. Analysis of the results showed that the behavior of single *Ltk^-/-^* or *Alk^-/-^* mutants was similar to that of WT mice at all ages tested (Fig. 4A). In the case of *DKO*s, mice displayed higher latency times for entry into the dark area and a significant reduction in transitions (Fig. 4A and Fig. S5A), indicative of reduced exploratory behaviour and increased anxiety that also became more pronounced with age.

**Figure 4.**
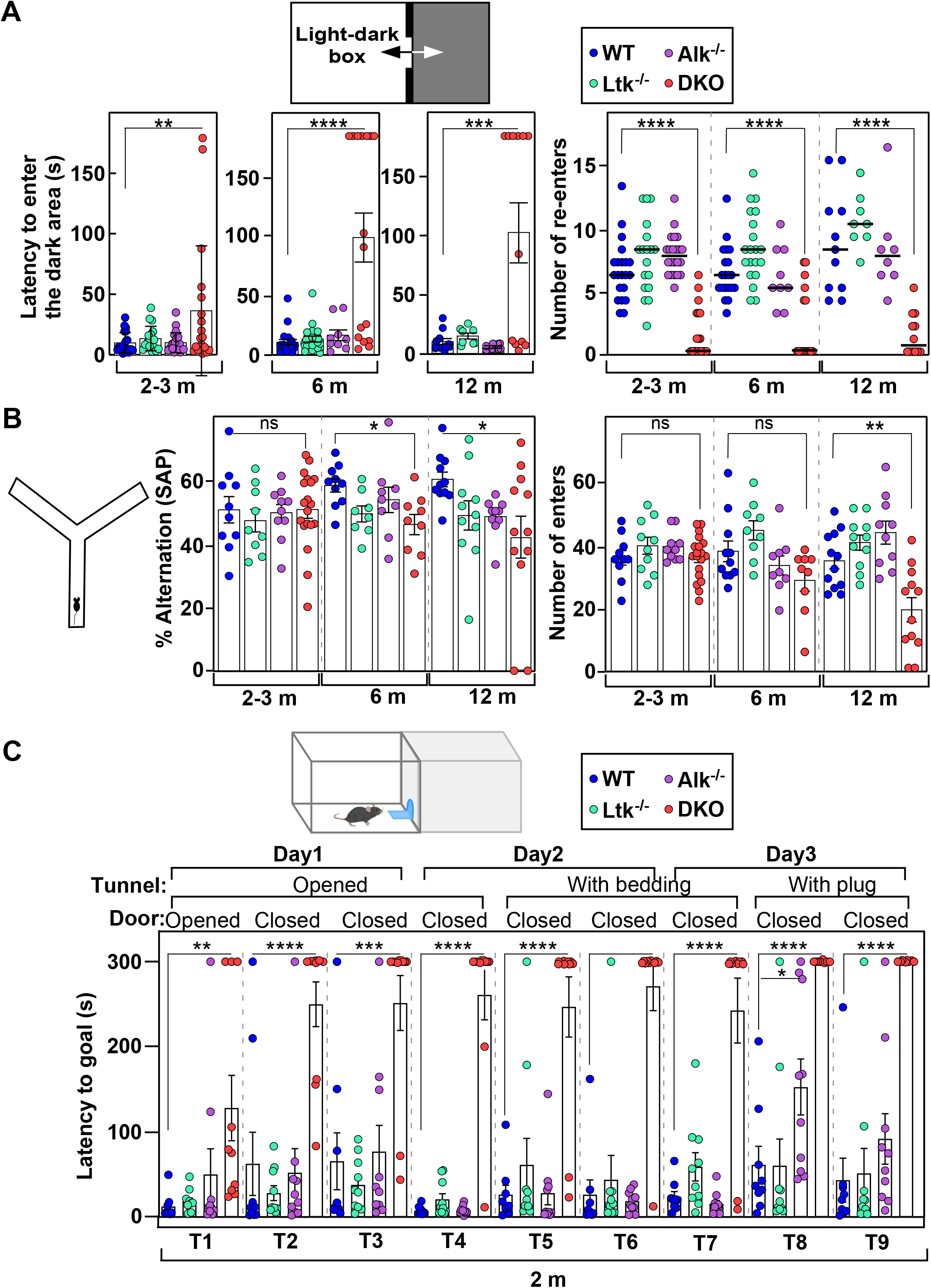
DKO mice exhibit behavior abnormalities. **(A)** Schematic of the light-dark box for assessing the anxiety levels. Latency to enter the dark box and the number of re-enters into the dark box for each individual mouse of the indicated genotypes and of various ages are plotted. **(B)** Schematic of the Y maze for assessing working memory. Spontaneous alternation (SAP) and number of arm entries for each individual mouse of the indicated genotypes are plotted. **(C)** Schematic of the puzzle box arena consisting of a white, light compartment (start box) and a black compartment (goal box) divided by black barrier. Latency to reach the goal box during the 9 trials of the test is plotted for each individual 2-3 m old mouse of the indicated genotypes. Data are plotted as mean ± SEM. Data were analyzed using one-way ANOVA and comparisons to WT were performed using Dunnett’s test (****p<0.0001, ***p<0.001, **p<0.01, *p<0.05, ns, not significant).

We next employed the Y-maze test for spontaneous alternations, which measures the tendency to make free choices in an alternating manner in the maze and is a paradigm for studying working memory (Kraeuter, Guest et al., 2019, Miedel, Patton et al., 2017). Mice were placed in one arm of the Y-maze and exploration of the three arms was monitored for 8 minutes by video tracking. The number of arms entered and the sequence of entries was recorded. A high alternation rate (degree of arm entries without repetitions) is indicative of sustained cognition as the animals must remember which arm was entered last to avoid re-entry. We found that spontaneous alternations (SAP) and the total number of arm entries by the single *Ltk^-/-^* or *Alk^-/-^* mice was comparable to wild-types at all ages tested. However, while young (2-3 month) *DKO*s performed similar to wild-type mice, a decrease was observed in older (6- and 12-month-old) *DKO*s (Fig. 4B). Thus, young adult mice of all genotypes display normal short-term spatial working memory, with some decline evident in older *DKO* mice.

Finally, to explore executive functions, the puzzle box assay was carried out (Ben Abdallah, Fuss et al., 2011). In this problem-solving test, mice are required to use a range of executive functions including working and contextual memory, spatial navigation and problem solving to complete escape tasks (T1-9) of increasing difficulty. Mice must move from a light area to a dimly lit goal box through a tunnel with barriers of increasing complexity introduced over a 3-day period. Repetition of tasks allows assessment of both short term (T3, T6, T9) and long-term memory (T4 and T7). Analysis of the results revealed that in young mice (2-3 months), *DKO*s displayed an increase in time (latency) of entry to the goal box for all tasks on all 3 days of the experiment as compared to single mutants or wild-type mice (Fig. 4C). The latency was notably increased upon the second challenge (T2) when the tunnel door was closed. In young adults (2-3 months), single *Ltk* or *Alk* knockouts showed behaviours similar to WT mice for all tasks, though performance in the most complex tasks (T8 and T9) significantly worsened with age (Fig. S5B and C). Thus, *DKO* mice showed severe deficits in problem-solving behaviour that also emerged in aging Ltk and Alk mutant mice for the most complex tasks. No effects on short- or long-term memory were observed in single KO mice, while it was not possible to assess this in *DKO*s who failed all tasks.

Taken together, these behavioural analyses demonstrate that while all genotypes display normal short-term memory, the single *Ltk^-/-^* or *Alk^-/-^* mice show age-related decline in the performance of complex functions while those lacking both Ltk and Alk, display severe anxiety and are unable to perform tasks requiring executive functions. Thus, loss of Ltk/Alk disrupts behavior and cognitive functions adult mice.

### The Alkal2 ligand activates Ltk/Alk to control neuronal polarity

We next sought to gain insights into the molecular basis for the disruption of neuronal polarity causes by loss of *Ltk/Alk*. The Ltk/Alk ligand, Alkal2, is expressed at roughly equivalent levels in both WT and *DKO* mice in the embryonic, newborn and adult brain and in isolated embryonic cortical neurons (Fig. S1G and Fig. S6A) indicating that changes in ligand expression do not explain the phenotype, thus, we next explored whether the ligand has a role in controlling neuronal morphology. Abrogating expression of Alkal2 using validated siRNAs yielded embryonic cortical neurons that displayed the multiple axon phenotype (Fig. 5A and Fig. S6B). Alkal2 protein is not commercially available, so to explore the effects of exogenously added ligand, we generated ALKAL2 conditioned media (ACM) from transiently transfected HEK293T cells as previously described (Guan et al., 2015). We first confirmed that the ALKAL2-conditioned media (ACM) was capable of activating LTK/ALK using IMR-32 human neuroblastoma cells that express wild-type ALK (Janoueix-Lerosey, Lequin et al., 2008) by immunoblotting with pY1586-ALK antibodies (Fig. S6C). Treatment of WT cortical neurons with ALKAL2-conditioned medium (ACM) inhibited the formation of axons (Fig. 5B) similar to the effect observed upon overexpressing LTK (Fig. 3C and Fig. S4B), but importantly, had no effect in murine *DKO* neurons (Fig. 5B). Thus, Alkal2 inhibits axon outgrowth in a manner dependent on the presence of endogenous Ltk/Alk receptors. Taken together, our results show that Alkal2-mediated activation of Alk and Ltk can regulate neuronal polarity.

**Figure 5.**
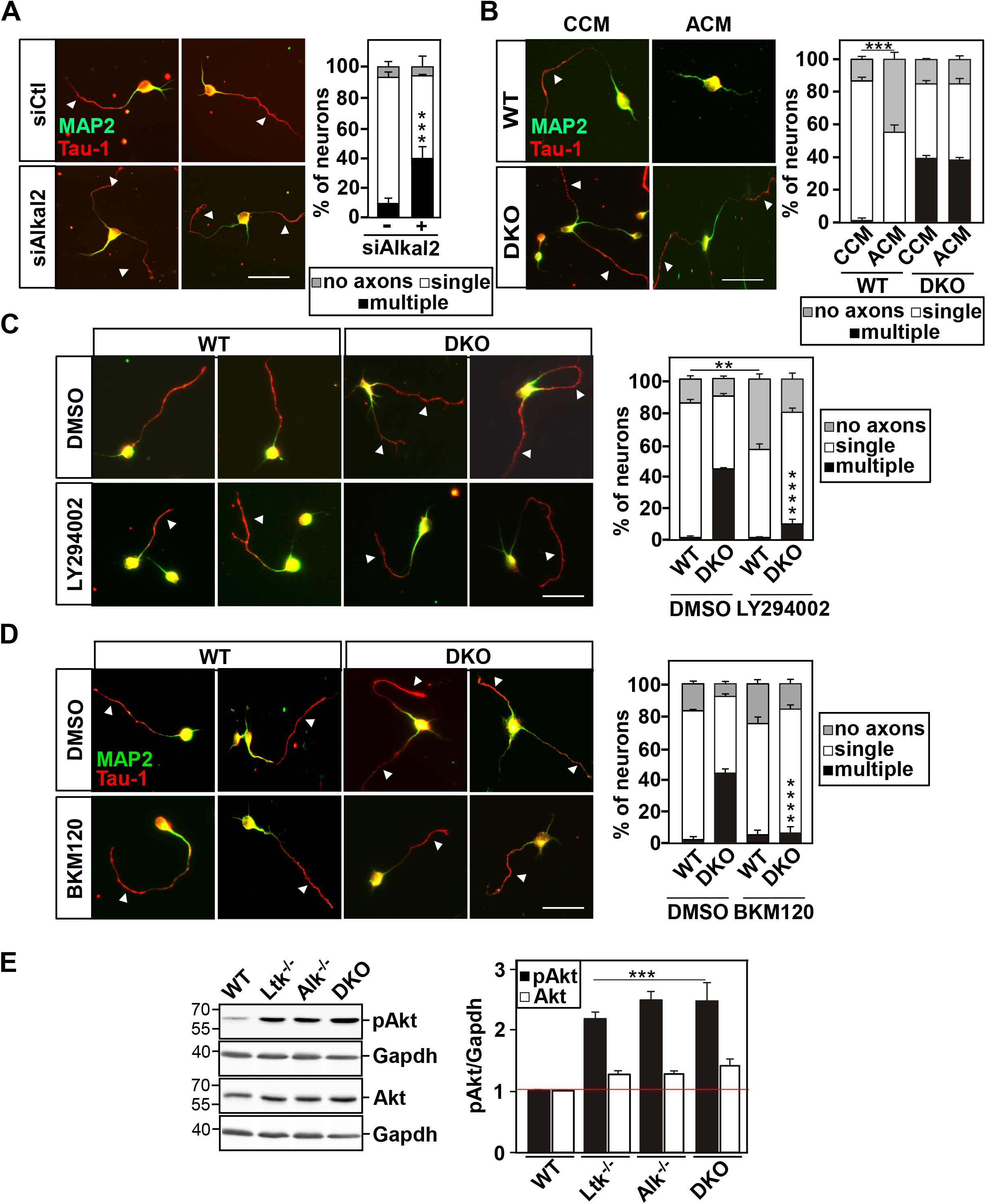
Analysis of the effects of ligand and inhibition of PI3 kinase activity on neuronal polarity. **(A)** RNAi-mediated depletion of Alkal2 in WT cortical neurons disrupts neuronal polarity. Dissociated E16 WT cortical neurons were transfected with siAlka2 or siCTL. **(B)** ALKAL2 ligand addition modulates neuronal polarity. WT and DKO cortical neurons were treated with control (CCM) or ALKAL2 (ACM) conditioned medium 4h after plating **(C, D)** Treatment of cortical neurons from WT or KO mice with PI3 kinase inhibitors reverses polarity defects caused by loss of Ltk/Alk. Dissociated E16 cortical neurons were treated with LY294002 (10 µM; **C**), BKM120 (1 µM; **D**) or DMSO control, 4 h after plating. (**A-D**) Representative images of cells fixed at 38h and stained with Tau-1 (axons, red) and MAP2 (dendrites, green) are shown. Arrowheads mark axons. Scale bars are 20 µm. Quantitation of the percent of neurons with multiple, single or no axons is plotted as the mean ± SEM of n = 150 neurons per condition from 3 independent experiments. **(E)** A representative immunoblot showing the levels of pAKT (pAkt^S473^), Akt and GAPDH (loading control) in lysates of cortical neurons from WT, single and DKO mice. Quantitation of pAkt^S473^ or Akt levels normalized to Gapdh relative to WT is plotted as the mean ± SEM of three independent experiments. Statistical significance: ****p<0.0001, ***p<0.001, **p<0.01 by one–way ANOVA, Dunnett’s test.

### Lack of Ltk/Alk expression results in excessive PI3K signalling

Phosphatidylinositol-3-kinase (PI3K) activity is a key driver of neuronal polarization as it is required and sufficient for axon specification (Yoshimura, Arimura et al., 2006), thus, we next sought to explore whether loss of Ltk/Alk engages the PI3K pathway. Inhibition of PI3K using the broad specificity inhibitor, LY294002 or with BKM120, a more selective pan-class I PI3K inhibitor that targets all isoforms of the p110 catalytic subunit of PI3K (Maira, Pecchi et al., 2012), blocked formation of axons in WT neurons (Fig. 5C and D). Remarkably, in neurons isolated from double KO mice, treatment with LY294002 or BKM120 not only prevented formation of excess axons, but restored the wild-type single axon phenotype (Fig. 5C and D). Similar results were observed in neurons isolated from single KO mice and in WT neurons in which the expression of Ltk, Alk or both was abrogated using siRNAs (Fig. S6D and E). Overexpression of the lipid and protein phosphatase, PTEN, which inactivates the PI3K pathway by dephosphorylating PIP3 (Jiang, Guo et al., 2005), also inhibited formation of axon-like extensions in wild-type cortical neurons and restored the single axon phenotype in *DKO* neurons (Fig. S6F).

Activation of PI3K results in increased phosphorylation of AKT. Thus, to confirm that the PI3K signalling pathway is activated in the absence of Ltk/Alk, lysates from isolated embryonic cortical neurons as well as from total brain homogenates from embryonic WT and KO mice were analyzed by immunoblotting using a phospho-specific Akt antibody. These results revealed that isolated neurons and brain extracts from single and/or double knockout mice display increased levels of pAKT (Fig. 5E and Fig. S6G). Thus, our results demonstrate that loss of Ltk/Alk results in excessive PI3K signaling that promotes the formation of multiple axons and that inhibition of PI3K activity can overcome the effects of loss of Ltk/Alk and restore the normal single axon phenotype in KO neurons.

### Igf-1r is activated in Ltk/Alk KO neurons

The observed increase in PI3 kinase activity in KO cortical neurons suggested the possibility that loss of Ltk/Alk may result in aberrant upstream activation of another receptor tyrosine kinase (RTK), as RTKs are well known activators of PI3K signalling. Insulin-like growth factor 1 receptor (IGF-1R) is widely expressed in the brain (Fernandez & Torres-Aleman, 2012) and in >90% of the isolated embryonic E15.5 cortical neurons used in our assays (Fig. S4A), and has been shown to play a role in neuronal polarity, migration and axonal outgrowth (Nieto Guil, Oksdath et al., 2017, O’Kusky & Ye, 2012, Sosa, Dupraz et al., 2006). Moreover, in ALK-positive lung cancer, acquisition of resistance to crizotinib is associated with enhanced IGF-1R activation, suggesting that IGF-1R may compensate for reduced ALK signaling during anticancer therapy (Lovly, McDonald et al., 2014). Thus, to test whether IGF-1R is activated in KO neurons, we immunoblotted neuronal lysates using a phospho-tyrosine (pY) antibody that recognizes the auto-phosphorylation sites in IGF-1R and the closely related Insulin receptor (InsR). This analysis revealed an increase in the levels of phosphorylated Y1135/1136 in cortical neurons in KO mice as compared to wild-type controls (Fig. S7A).

To further explore whether IGF-1R and/or InsR are activated upon loss of Ltk/Alk, we first tested the effect of an inhibitor, tyrphostin, AG1024, which inhibits IGF-1R with 8-fold higher affinity than InsR (ie 7 µM vs 57 µM, respectively (Parrizas, Gazit et al., 1997), and acts by preventing binding of substrate and ATP. Treatment of cortical neurons with AG1024 at a dose (1 µM) at which only IGF-1R but not InsR was inhibited restored the single axon phenotype in *DKO* neurons (Fig. 6A). Similarly, the small toxin molecule, picropodophylin (PPP), which specifically inhibits IGF-1R, but not InsR by blocking phosphorylation of Y1136 and receptor activation (Vasilcanu, Girnita et al., 2004), caused a marked decrease in the number of neurons with multiple axons in both single and double *DKO* cells (Fig. 6B and C and Fig. S7B). Consistent with these observations, treatment of neurons with a neutralizing anti-IGF-1Rα antibody, which prevents binding of the IGF-1 ligand to IGF-1R and thus blocks IGF-1-mediated receptor activation, revealed a restoration of the single axon phenotype in *DKO* neurons (Fig. 6C). Finally, siRNA-mediated abrogation of Igf-1r expression in *DKO* cortical neurons switched the multiple axon phenotype into the normal single axon morphology (Fig. 6D and E). Of note, in WT neurons, inhibition of Igf-1r receptor activity or silencing of Igf-1r increased the number of neurons with no axons (Fig. 6A, B and D), consistent with previous observations demonstrating a role for Igf1-r in neuronal polarity (Nieto Guil et al., 2017, Sosa et al., 2006). Finally, we examined the effect of inhibiting the Igf1-r on neuronal migration *in vivo*. Remarkably, treatment of DKO embryos with PPP led to a partial restoration of neuronal migration in P2 mice, with some BrdU+ neurons being localized in the superficial layers (Bin 7 and 8) (Fig. 6F). Of note, in WT mice, PPP treatment blocked migration of BrdU labelled neurons (Fig. 6F), indicating that Igf-1r activity is required for neuronal migration as previously reported (Nieto Guil et al., 2017). Thus, taken together our results indicate that Igf-1r activity is enhanced upon loss of Ltk/Alk and this can drive the multiple axon phenotype *in vitro* and drive defects in neuronal migration *in vivo*.

**Figure 6.**
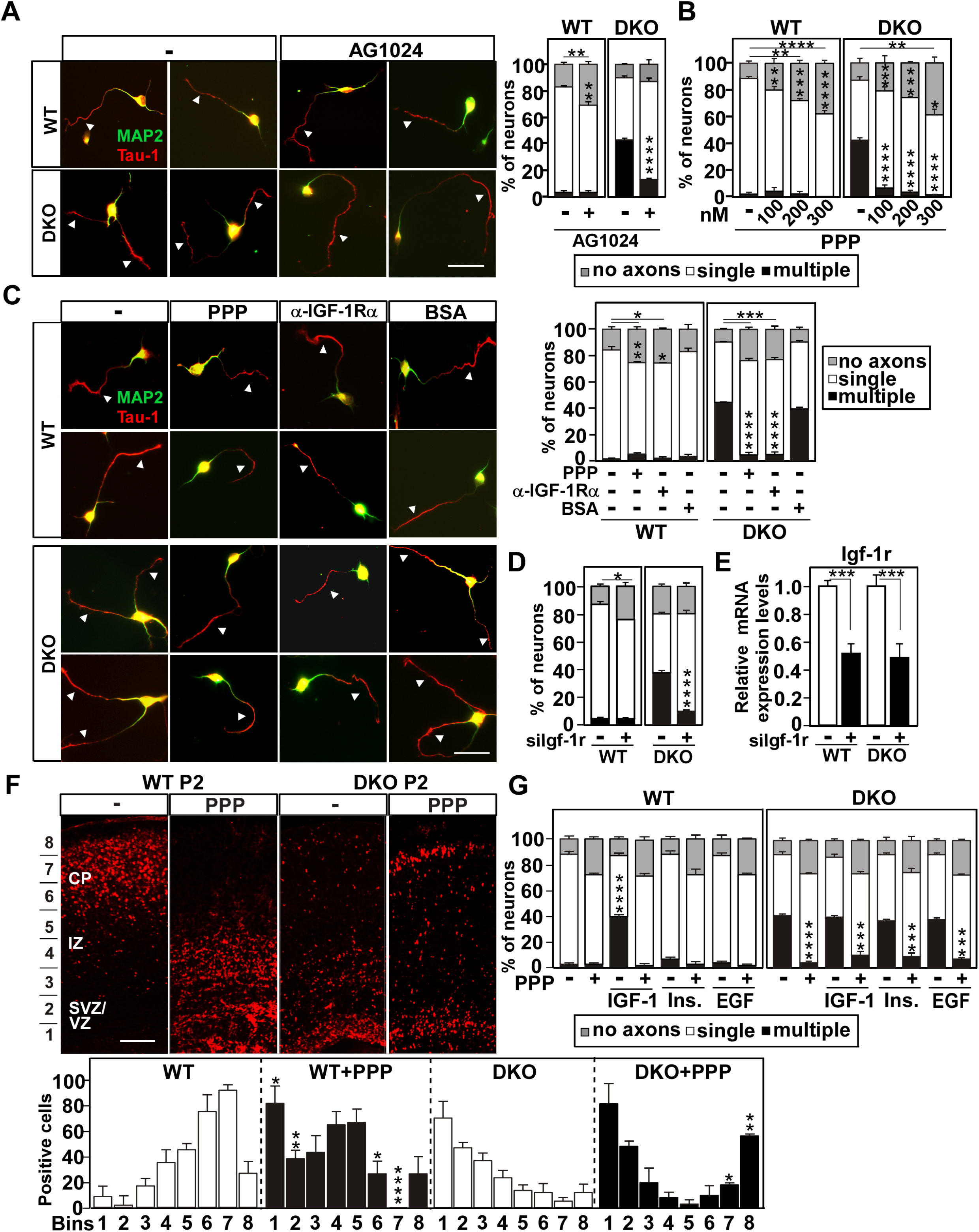
Loss of Ltk/Alk promotes Igf-1r activity. **(A, B)** IGF-1R inhibitors rescue the multiple axon phenotype in DKO cortical neurons. Cortical neurons, isolated from WT and DKO mice were treated with 1 µM AG1024 **(A),** varying concentrations of Picropodophylin (PPP; **B**) or DMSO as control 4 h after plating, fixed at 38 h, and stained for Tau-1 (axons, red) and MAP2 (dendrites, green). Quantitation of the percent of neurons with multiple, single or no axons is plotted as the mean ± SEM, n = 150 neurons from 3 independent experiments. **(C)** Blocking Igf-1r activation with a neutralizing anti-IGF-1Rα antibody rescued the multiple axon phenotype in DKO cortical neurons. Neurons isolated from WT or DKO mice were treated anti-IGF-1Rα antibody (1.5 µg/ml), BSA as control (1.5 µg/ml), or PPP (200 nM) 4 h after plating. Representative images of Tau-1 and MAP2 stained cells with arrowheads marking axons are shown. Scale bars are 20 µm. Quantitation of the percent of neurons with multiple, single or no axons is plotted as the mean ± SEM of n = >150 neurons per condition from 3 independent experiments. **(D)** Reduced expression of Igf-1r restores normal neuronal morphology in DKO cortical neurons. Cortical neurons isolated from WT and DKO mice were electroporated with siIgf-1r together with a GFP plasmid to mark transfected cells. Neurons were fixed at 36 h and stained for Tau-1 and MAP2. Quantitation of the percent of neurons with multiple, single and no axons in EGFP positive neurons is plotted as the mean ± SEM (n=150 neurons) from 3 independent experiments. **(E)** Knockdown efficiency was confirmed by qPCR. **(F)** Inhibition of Igf-1r blocks neuronal migration *in vivo*. Pregnant E14 WT or DKO mice were injected (i.p) with BrdU and 24 h later were injected with PPP or saline as control. Brains from P2 pups were subjected to staining with anti-BrdU antibody. Quantitation in P2 cortices of BrdU-positive cells in the indicated bins (marked on left) is plotted as the mean +/- SEM from 3 (DKO) or 2 (WT) independent experiments. Statistical significance: ****p<0.0001, ** p<0.01, *p<0.05 by Student’s t-test where WT vs WT+PPP or DKO vs DKO+PPP were compared. The location of the cortical plate (CP), intermediate zone (IZ), subventricular/ventricular zone SVZ/VZ in P2 mice and are indicated. Scale bars are 50 µm. **(G)** Analysis of the effect of ectopically applied recombinant IGF-1. Cortical neurons isolated from WT or DKO mice were treated with PPP (200 nM), DMSO as control and /or IGF-1 (50 nM), insulin (100 nM) or EGF (20 ng/ml). Cells were fixed at 38h and stained for Tau-1 and MAP2. Quantitation of the percent of neurons with multiple, single or no axons is plotted as the mean ± SEM, n=100 neurons from 3 independent experiments. **(A-E,** and **G)** Statistical significance: ****p<0.0001, ***p<0.001, **p<0.01, *p<0.05. by one–way ANOVA, Dunnett’s test

Given that loss of Ltk/Alk enhances Igf-1r activity, we next examined the effect of activating Igf-1r in the context of wild-type neurons by ectopically adding murine, recombinant IGF-1. Cortical neurons treated with IGF-1 formed multiple axons in a dose dependent manner (Fig. S7C), with the effect of 50 nM IGF-1 being similar to that observed for untreated *DKO* cortical neurons (Fig. 5G and Fig. S7C). In contrast, neither insulin (a ligand for InsR/IGF-1R) nor EGF (a ligand for EGFR) altered the number of axons and axon-like neurites (Fig. 6G and S7C). The effect of IGF-1 was completely abolished by pre-treatment of cells with the IGF-1R inhibitor, PPP, consistent with the notion that activation of Igf-1r leads to the formation of multiple axons (Fig. 6G). In *DKO* cortical neurons addition of IGF-1, however, did not increase further the number of neurons bearing multiple axons, suggesting that Igf-1r activity is already saturated in *DKO* neurons (Fig. 6G and Fig. S7C). Altogether, these data provide compelling evidence that Igf-1r is activated in cortical neurons upon loss of Ltk/Alk and that this increases PI3 kinase activity to promote the formation of multiple axons. Our results also show that pharmacologically blocking IGF-1R activity or abrogating Igf-1r expression can restore the normal single axon phenotype in KO cortical neurons.

### Ltk/Alk modulates Igf-1r levels and activity

We next conducted studies to explore the molecular mechanism that would lead to increased Igf-1r activity in KO neurons. Analysis of Igf-1r mRNA expression by real-time PCR revealed no differences in WT versus KO cortical neurons (Fig. S7D) indicating a post-transcriptional mechanism. We next examined Igf-1r protein levels and activity by immunoblotting using antibodies that recognize Igf-1r total protein or the phosphorylated (pY1135/1136), activated Igf-1r/InsR. This revealed that *DKO* cortical neurons display a modest increase in total Igf-1r protein levels (<2 fold) along with a more dramatic enhancement in the levels of activated Igf-1r as compared to WT controls even when corrected for the increase in protein levels (Fig. 7A). Thus, the increase in Igf-1r activity in KO cortical neurons is not a result of transcriptional regulation but rather due to post-transcriptional enhancement of the levels of both total protein and activated receptors.

**Figure 7.**
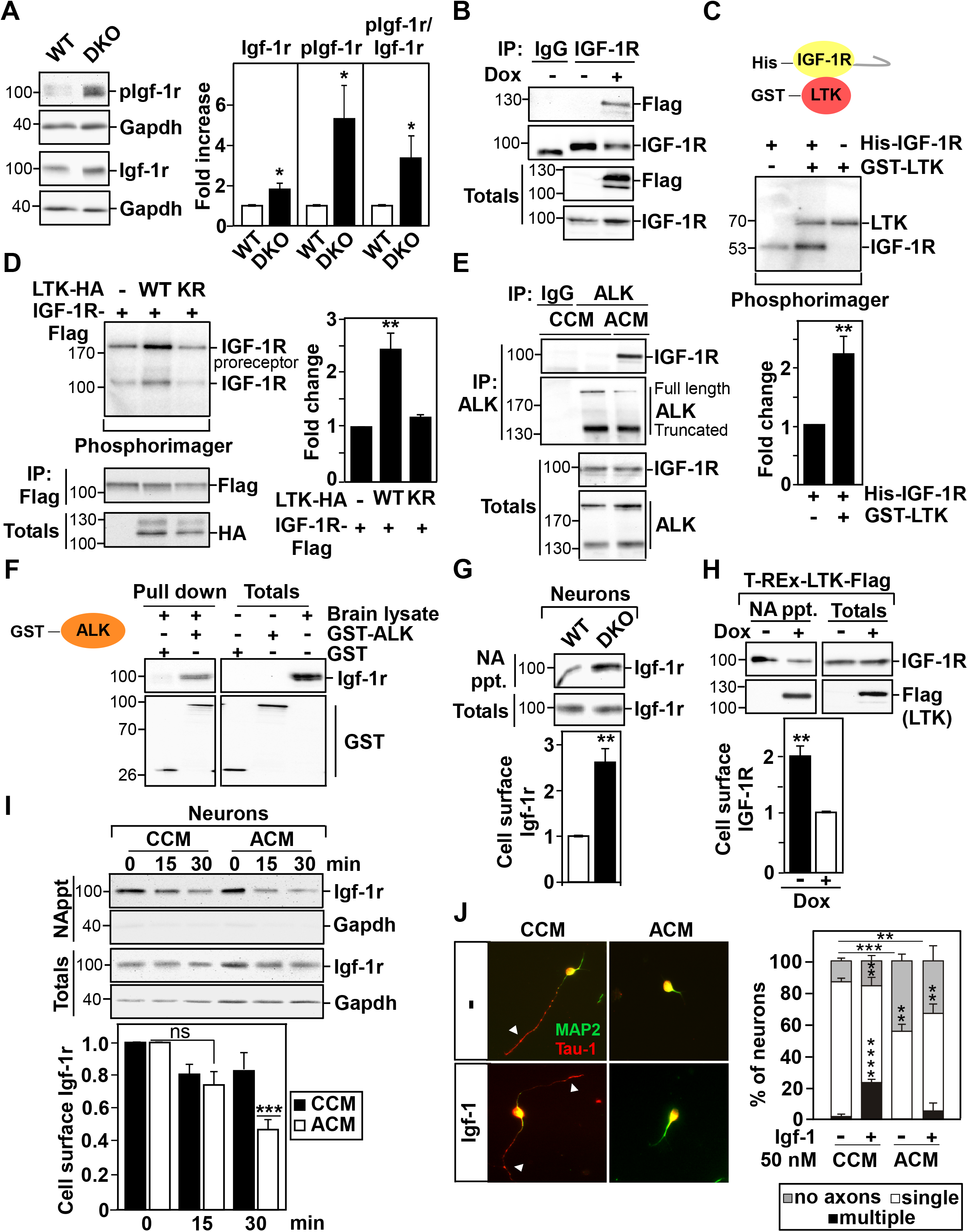
Ltk/Alk interacts with and modulates Igf-1r stability and cell surface expression. **(A)** Representative immunoblot showing the levels of phosphorylated (pIgf-1r^Y1135/1136^), total Igf-1r and Gapdh (as loading control) in lysates of isolated cortical neurons from WT and DKO mice. Bands intensities for pIgf-1r and Igf-1r were normalized to Gapdh and the fold increase relative to WT is plotted as the mean ± SEM of 3 independent experiments. **(B)** Endogenous IGF-1R was immunoprecipitated from cell lysates of T-REx cells, treated without (-) or with (+) doxycycline (Dox) to induce expression of Flag-tagged LTK and the interaction with LTK-Flag was visualized by immunoblotting. **(C)** Representative image showing phosphorylation of recombinant His-tagged IGF-1R (kinase domain and C-terminal region) by recombinant GST-LTK (kinase domain) in an *in vitro* radioactive kinase assay. Phosphorylation of IGF-1R was quantitated as fold change relative to auto-phosphorylation of IGF-1R alone and plotted as mean ± SEM from 7 independent experiments. **(D)** Representative image showing the phosphorylation of IGF-1R by co-expressed WT or Kinase-deficient (KR) LTK in *in vitro* radioactive kinase assay. The levels of immunoprecipitated IGF-1R and total LTK-HA were visualized by immunobloting (bottom). Phosphorylation of IGF-1R was quantitated as fold change relative to auto-phosphorylation of IGF-1R alone and plotted as mean ± SEM from 5 independent experiments. **(E)** Endogenous ALK was immunoprecipitated from lysates of IMR-32 cells, treated with control (CCM) or ALKAL2 (ACM) conditioned media and co-precipitated endogenous IGF-1R was detected by immunoblotting. **(F)** Homogenates from WT embryonic brains were incubated with GST-ALK or GST alone, immobilized to GSH-Sepharose beads. Bound and total proteins were subjected to immunoblotting with anti-IGF-1R antibody. **(G)** Cell-surface expression of Igf-1r is increased in DKO cortical neurons. Cell-surface Igf-1r (NA ppt) is normalized to totals and the fold change relative to WT samples is plotted as the mean ± SEM from three independent experiments. **(H)** LTK decreases cell-surface expression of IGF-1R in T-Rex-LTK stable cells. Cell-surface IGF-1R (NA ppt) normalized to totals as fold change relative to samples expressing LTK (+Dox) is plotted as the mean ± SEM from 3 independent experiments. **(I)** WT cortical neurons were labeled with biotin and incubated at 37°C for 15 or 30 min in the presence of CCM or ACM conditioned media. Cell-surface Igf-1r (NA ppt) normalized to totals as the fold change relative to controls at time 0 is plotted as the mean ± SEM from 4 independent experiments. **(J)** WT and DKO cortical neurons were treated with control (CCM) or ALKAL2 (ACM) conditioned medium 4h after plating, and IGF-1 was added 1 hour later. Representative images of cells fixed at 36 h and stained for Tau-1 (axons; red) and MAP2 (dendrites: green) are shown. Arrowheads mark axons. Quantitation of the percent of neurons with multiple, single or no axons is plotted as the mean ± SEM of n = 150 neurons per condition from 3 independent experiments. **(A-B, E-G)** Representative blots from 3 independent experiments are shown. Statistical significance: ****p<0.0001, ***p<0.001; **p<0.01; *p<0.05 by Student’s t-test.

To determine how Ltk/Alk might modulate Igf-1r, we explored whether Ltk/Alk form a complex with Igf-1r. When transiently overexpressed in HEK293T cells, an association between IGF-1R and either LTK or ALK was detected (Fig. S7E and F). In LTK expressing T-REx stable cells, the interaction of endogenous IGF-1R with Dox-induced Flag-tagged LTK was also observed (Fig. 7B). This interaction was substantially reduced when a kinase-inactive variant (KR) of LTK was used, indicating that the association requires active LTK (Fig. S7G). In contrast, IGF-1R kinase activity was unnecessary for efficient interaction with LTK (Fig. S7H). To examine whether LTK/ALK might phosphorylate IGF-1R, we performed an *in vit*ro kinase assay using recombinant proteins, comprised of the kinase domain alone of LTK (GST-LTK) and the kinase and C-terminal tail of IGF-1R (His-IGF-1R). Addition of GST-LTK increased the level of phosphorylation on IGF-1R over the basal levels of IGF-1R autophosphorylation, indicating that LTK can phosphorylate IGF-1R (Fig. 7C). A GST pull-down assay confirmed that the kinase domain of LTK was sufficient for interaction with the kinase/C-terminal region of IGF-1R (Fig. S7I). Next, we analyzed IGF-1R phosphorylation in HEK293T cells transiently-transfected with Flag-tagged IGF-1R with or without HA-tagged WT or KR variants of LTK. IGF-1R was immunoprecipitated and subjected to an *in vitro* kinase assay. Consistent with the assay using recombinant proteins, IGF-1R is phosphorylated by WT but not kinase inactive LTK (Fig. 7D). IMR-32 cells express both endogenous ALK and IGF-1R, thus we examined the effect of ligand on receptor interactions. Immunoprecipitation of ALK revealed an association with IGF-1R that was dependent on treatment with ALKAL2 (ACM) conditioned media, confirming the importance of ligand-induced activation of ALK for the interaction with IGF-1R (Fig. 7E). Finally, pulldown assays from embryonic mouse brain lysates using bacterially expressed recombinant GST-tagged kinase domains of active ALK or LTK showed efficient isolation of endogenous Igf-1r (Fig. 7F and Fig. S7J), confirming the association and indicating that the kinase domain of LTK/ALK is sufficient for this interaction. Taken together, these findings support the notion that active LTK/ALK interacts with and phosphorylates IGF-1R.

Given the increased level of active Igf-1r and since receptors signal from the plasma membrane, we next examined whether the presence of Ltk/Alk might modulate the levels of Igf-1r on the cell-surface. For this, cell-surface proteins were biotinylated, cells were lysed, biotin-labelled proteins were collected with NeutrAvidin-Sepharose beads and bound proteins were then detected by immunoblotting. In dissociated cortical neurons, we observed that the levels of cell-surface localized Igf-1r were significantly increased in neurons from *DKO* mice when compared to WT neurons (Fig. 7G). In the converse experiment using T-REx cell stably expressing doxycycline (DOX)-inducible LTK, we observed that increasing LTK expression resulted in a decrease in cell-surface localization of endogenous IGF-1R (Fig. 7H). Thus, LTK/ALK attenuates cell-surface expression of IGF-1R and accordingly loss of Ltk/Alk enhances cell-surface localization of Igf-1r. We next examined the effect of Ltk/Alk receptor activation on Igf-1r cell-surface expression and protein levels in WT neurons. For this, cell-surface receptors on cortical neurons were labeled with biotin, and cells were then incubated at 37°C for varying times with control (CCM) or ALKAL2 (ACM) conditioned media to activate Ltk/Alk. Remarkably, after 30 min of treatment with ACM, a 50% reduction in Igf-1r on the cell-surface was observed, indicating a more rapid internalization of Igf-1r in the presence of active Ltk/Alk (Fig. 7I). Consistent with the observed decrease in cell surface Igf-1r upon activation of Ltk/Alk, addition of Igf-1 had no effect on axon formation in neurons in which Ltk/Alk receptors had been previously activated by ALKAL2 (Fig. 7J). Thus, ligand-induced activation of Ltk/Alk favors Igf-1r internalization and results in a decrease in Igf-1r cell-surface expression.

Taken together, our results reveal a key role for Ltk/Alk in controlling neuronal morphology and patterning of the cortex and provide a molecular model for how Ltk/Alk functions in this context (Fig. 8). Specifically, Alkal2-activated Ltk/Alk act to limit the level of cell-surface, activated Igf-1r by promoting Igf-1r internalization to maintain proper neuronal morphology. Loss of Ltk/Alk increases Igf-1r cell surface localization and activity, to enhance PI3K signalling and thereby promotes the formation of multiple axons. *In vivo*, loss of Ltk/Alk disrupts neuronal polarization, delays neuronal migration and cortical patterning and in adults leads to an aberrant morphology of cortical neurons and disrupts axon tracts in the corpus callosum. Altogether, this highlights a key role for Ltk/Alk in cortical development and patterning.

**Figure 8.**
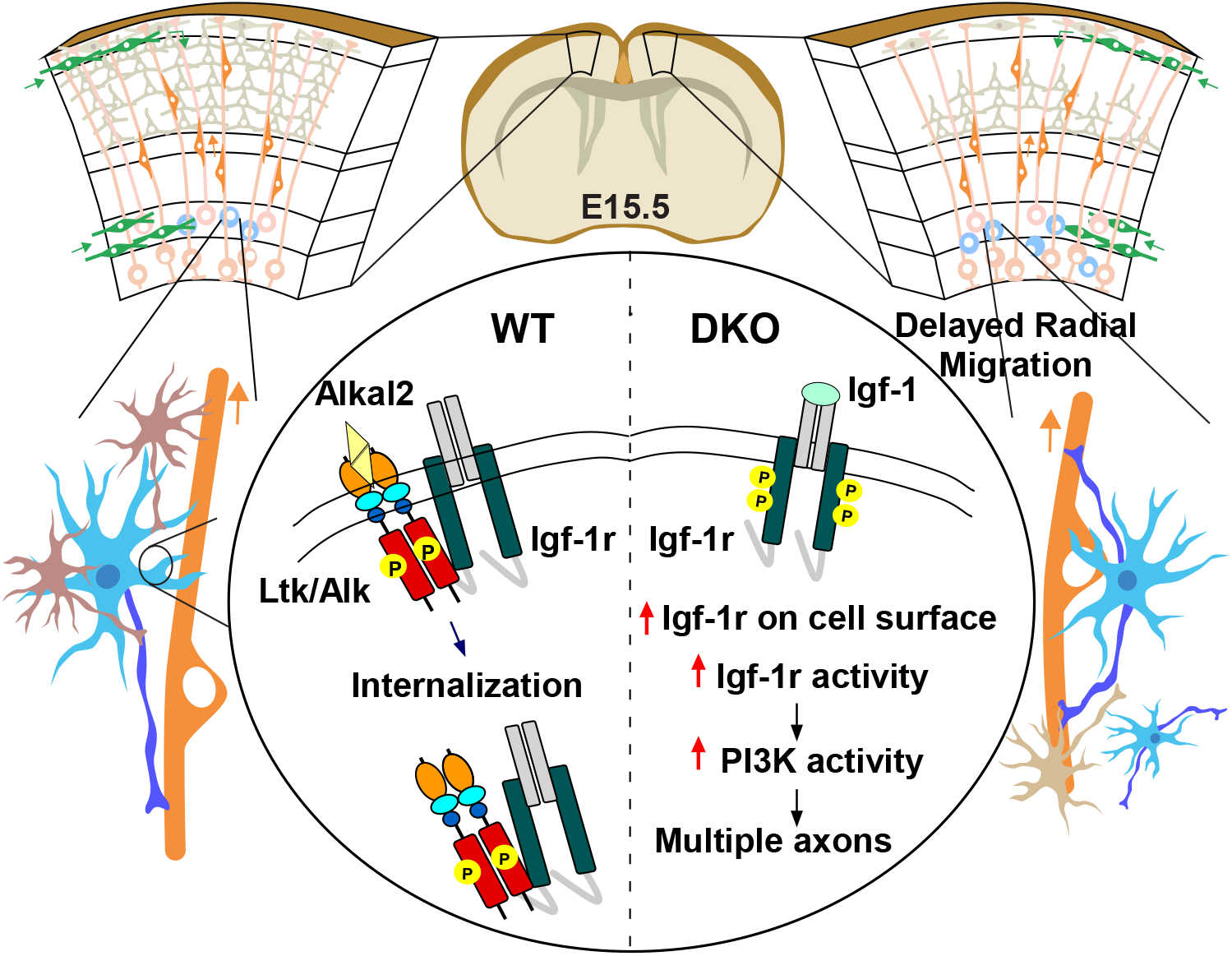
A model depicting the role of Ltk and Alk in regulating neuronal morphology through the modulation of Igf-1r activity.

## Discussion

The establishment of axon-dendrite polarity is fundamental for radial migration of neurons during mammalian cortical development and subsequently for proper formation of the neuronal circuitry. Here, we demonstrate that the receptor tyrosine kinases, Ltk and Alk, contribute to polarization of developing neurons and the subsequent timing of cortical patterning. We observed that Ltk, Alk and the Ltk/Alk ligand, Alkal2 are expressed in the developing cortex, including in the embryonic progenitor cell populations found in the ventricular and subventricular zones, where neuronal polarization occurs. Cortical neurons exhibit the highest migratory activity between E14.5 to E16.5 which also continues in newborn pups, up to P7 (Chen, Sima et al., 2008). Proper neuronal morphology is essential for neuronal migration and in line with this, we observed that in Ltk and Alk single or double knockouts (*DKOs)*, the radially migrating cortical neurons in E17.5 and newborn pups display a delay in localization to the correct position within the defined layers of the developing cortex.

Consistent with the *in vivo* observations, our analysis of isolated primary embryonic neurons confirmed that loss or inhibition of Ltk/Alk activity using siRNAs or small molecule inhibitors or in embryonic neurons extracted from KO mice, there are defects in neuronal polarization. These neurons display the presence of multiple axons, a phenotype that was rescued by overexpression of Ltk/Alk. Of the two known and related Ltk/Alk ligands (Reshetnyak et al., 2015), only Alkal2 was expressed in the developing and adult cortex. Consistently, a deficiency in Alkal2 expression or activation of Ltk/Alk receptors by the addition of exogenous ALKAL2, yielded neuronal phenotypes resembling those of abrogated or overexpressed Ltk/Alk receptors, respectively, thereby highlighting the functional link of ligand-mediated receptor activation in the regulation of neuronal morphology. During the course of our experiments, we observed that loss of either Ltk or Alk alone or both together yielded a similar proportion of isolated neurons with a multiple axon phenotype. Our single-molecule RNA *in situ* analysis demonstrated that the bulk (∼70%) of isolated neurons express both Ltk and Alk as well as Igf-1r. These results, thus, indicate that in embryonic neurons Ltk or Alk cannot compensate for loss of the other, rather that both Ltk and Alk are required to regulate neuronal polarity. We made similar observations *in vivo,* where loss of either Ltk or Alk alone or both together, resulted in defects in neuronal migration and cortical patterning. Taken together, these data lead us to speculate that Ltk and Alk may act together in the context of a heteromeric Ltk/Alk complex to regulate neuronal polarity (Fig. 8). While recent structural insights into ligand mediated receptor activation have focused on ALK or LTK homodimers (Reshetnyak et al., 2021), the formation of heteromeric Ltk/Alk complexes in the presence of a ligand has been previously reported (Guan et al., 2015).

The mouse brain continues to develop postnatally and interestingly, the impaired cortical layering present at birth, showed improvements by P7, and appeared resolved by adulthood. However, it is worth noting that despite the apparent recovery of normal cortical patterning, adult mice display neurons with morphological defects including the presence of aberrant neuronal projections within the cortex. In addition, there is a disruption of the axon tracts within the corpus callosum, which are derived primarily from neurons from cortical layers II-V. While we have not directly demonstrated that the defects in the axon tracts is the result of excess Igf1-r activity, it is known that defects in the corpus callosum can arise from earlier disruptions in neuronal polarization (Edwards, Sherr et al., 2014), indicating that the aberrant axon tracts may be the result of effects on neuronal morphology. Consistent with the presence of alterations in adults, *DKO* mice display behavioral anomalies including severe anxiety and the inability to perform complex tasks requiring executive functions. In general, the single *Ltk^-/-^* or *Alk^-/-^* mice behaved similar to WT though an age-related decline in the performance of complex functions was observed. This suggests that in single mutant adults, compensatory mechanisms such as re-wiring of neuronal circuits can overcome the severity of the defect. It is also possible that there are Igf1-r independent, but redundant Ltk/Alk pathways that result in the behavioural defects only observed in the DKOs, such as the recently reported ALK-LIMK-cofilin pathway which regulates synaptic scaling (Zhou, He et al., 2021). It is worth noting that in mice deficient in Alk alone, others have reported subtle behavioural alterations, altered responses to ethanol and hypogonadotropic hypogonadism (Bilsland et al., 2008, Lasek et al., 2011, Weiss et al., 2012, Witek et al., 2015).

Our mechanistic investigations in cultured neurons revealed that the presence of additional axons was due to excess PI3 kinase activity resulting from activation of Igf-1r signalling that occurred upon the loss of Ltk/Alk (Fig. 8). Consistent with these studies, administration of a small molecule inhibitor of Igf-1r *in vivo* partially rescued the neuronal migration defects observed in Ltk/Alk knockout mice indicating excess Igf-1r activity also contributes to the migrational delay. IGF-1R belongs to the subfamily of insulin transmembrane receptor tyrosine kinases, is widely expressed in the brain and has a well-studied role in neuronal polarity, migration and axonal outgrowth (Nieto Guil et al., 2017, O’Kusky & Ye, 2012, Sosa et al., 2006). Activation of IGF-1R engages the PI3 kinase signalling pathway, which leads to the accumulation of PIP3 in the growth cone that in turn is critical for regulating membrane expansion and axonal outgrowth (Laurino, Wang et al., 2005, Sosa et al., 2006). In line with this, we showed that increased PI3K activity as a result of the loss of Ltk/Alk could be attributed to an increase in Igf-1r cell-surface localization and activity. Ligand-induced activation of Ltk/Alk in WT neurons resulted in a loss of biotin-labelled cell-surface Igf-1r, which is strongly indicative of increased internalization. Since internalization can lead to receptor degradation, this is consistent with our observation of a post-transcriptional increase in Igf-1r protein levels upon loss of Ltk/Alk. It is worth noting that changes in cell-surface expression of Igf-1r, have also been reported to occur through exocytosis and that this process is essential for membrane expansion during axon formation (Quiroga, Bisbal et al., 2018). Thus, the increase in cell-surface expression of Igf1-r in DKO neurons may also be explained by enhanced exocytosis. In either case, the consequent multi-axon phenotype could be prevented using pharmacological inhibitors of Igf-1r, siRNAs targeting Igf-1r or neutralizing anti-IGF-1Rα antibodies. The compensatory upregulation of Igf-1r upon loss of Ltk/Alk activity is strikingly similar to reports that chronic inhibition of ALK in non-small cell lung cancer (NSCLC) patients leads to upregulation of Igf1-r activity (Lovly et al., 2014). Our studies further revealed that Ltk/Alk interact with Igf-1r in a manner that is dependent on Ltk/Alk activation, that Ltk/Alk can promote phosphorylation of Igf-1r and that this interaction results in a decrease in cell-surface expression and signalling of Igf-1r. RTKs are subject to numerous phosphorylation events, some of which are critical for receptor trafficking. In the case of Igf-1r, identifying and characterizing these events is an ongoing, active field of investigation (Rieger & O’Connor, 2020), with a recent study reporting that Y1250 phosphorylation promotes Igf-1r internalization (Rieger, O’Shea et al., 2020). Thus, the precise details of how Ltk/Alk mediated phosphorylation of Igf-1r promotes loss of cell-surface Igf-1r remains to be determined.

Altogether, our work uncovers a new role for Ltk/Alk as regulators of neuronal polarity and migration and indicates that correct neuronal morphology requires tightly regulated activity and expression of Ltk/Alk and their ligand. Aberrant activation of ALK due to protein fusions, point mutations, and overexpression have been described in many cancers (Hallberg & Palmer, 2016, Palmer et al., 2009) and a recent study provided compelling evidence that overexpression of the ligand, ALKAL2 in mice can drive neuroblastoma (Borenas et al., 2021). ALK inhibitors are in use in the clinic for the treatment of a variety of ALK-driven tumors, including NSCLC, primarily occurring as aberrant fusion proteins since ALK/LTK are not normally expressed in the lung. Second and third generation ALK inhibitors, such as ceritinib, alectinib, brigatinib and lorlatinib have been designed to cross the blood brain barrier as a means to treat the frequently observed brain metastasis in ALK-driven NSCLC tumors treated with crizotinib (Awad & Shaw, 2014, Chuang & Neal, 2015). Interestingly, a phase 2 clinical trial for lorlatinib showed that about 39% of patients (of 275 in the trial) displayed CNS effects that included changes in cognitive function, mood and speech (Solomon et al., 2018). Thus, our study could provide mechanistic insights into the basis for these adverse effects and further suggests that co-treatment with IGF-1R inhibitors may be a route for mitigating these effects. As the development of resistance in tumors to virtually all kinase inhibitors is inevitable, our study also provides a potential therapy whereby co-treatment with IGF-1R inhibitors may impact outcome. Our findings may also raise awareness for the use of ALK inhibitors for cancer treatment of females during pregnancy or lactation or in infant and young cancer patients and suggest that more considerations should be given to the potential risk of adverse complications during brain development.

## Materials and Methods

### Animals

All animal experiments were performed in accordance with the Animal Care and Use Committee of the University of Toronto. Mice were group housed (up to 4 per cage) with temperature controlled at 21-22°C and with 12-hour light-dark cycles. Animals were given food and water *ad libitum*. Wild-type CD-1 and C57BL/6 mice were purchased from Charles River and mutant animals were bred in house. *Alk^-/-^* and *Ltk^-/-^* knockout mice were generous gifts from Dr. Stephen Morris (St-Jude CRH, Memphis) and Dr. Joseph Weiss (OHSU, Oregon), respectively. *Alk^-/-^Ltk^-/-^* double knockout animals were generated by crossing homozygous *Alk^-/-^* and *Ltk^-/-^* single knockout mice. Genotyping was performed by PCR analysis using primers listed in Table S1. Mutant *Ltk* and *Alk* alleles produced a band of 400 base pairs in PCR and the wild-type allele produced a 500 base pair product.

### Antibodies, reagents, cDNA constructs and siRNAs

All primary antibodies used in the study are listed in Table S3. The following secondary antibodies and dyes were used: Alexa Fluor (488, 546, 594, 647, goat anti-mouse IgG1 cross-adsorbed Alexa Fluor 488 or 350 and anti-mouse IgG2a cross-adsorbed Alexa Fluor-568), Phalloidin-Rhodamine and 4’, 6-diamidino-2-phenylindole (DAPI) from ThermoFisher; anti-mouse IgG-horse radish peroxidase (HRP), anti-rabbit IgG-HRP and anti-rat IgG-HRP from Jackson Laboratories. The pcDNA3-FAM150B-HA plasmid, encoding ALKAL2 (FAM150B) (Guan et al., 2015) was a gift from Ruth Palmer (University of Gothenburg, Sweden). LTK, ALK and IGF-1R constructs were generated using the Gateway system (Life Technologies) by transferring from Gateway entry clone vector into C-terminally tagged pCMV5-based destination vectors. LTK-K544R and IGF-1R-K1003R mutants were generated by PCR-mediated site directed mutagenesis. For gene silencing experiments, siRNA for Ltk, Alk, Igf-1r, Alkal2 and RNAi negative control were purchased from Dharmacon (siCTL-ON TARGET plus Non targeting siRNA #3 D-001810-03; siAlk-siGenome Mouse Alk (111682) siRNA, set of 4, MQ-0401104-01; siLtk-siGenome Mouse Ltk, D-063855-04; siIgf-1r-siGenome mouse Igf-1r (16001), set of 4, MQ-056843-01; siAlkal2-siGenome mouse Fam150b (100294583) siRNA, set of 4, MQ-042520-00).

### Cell culture and transfections

Human embryonic kidney cells HEK 293T were obtained from American Type Culture Collection (ATCC) and cultured in DMEM supplemented with 10% fetal bovine serum (FBS, Gibco). T-REx-LTK stable cells, expressing Doxycycline-inducible Flag-tagged LTK, were generated by Gateway cloning (Invitrogen) and were grown in DMEM supplemented with 10% FBS, 1% penicillin/streptomycin (Gibco), 200 µg/ml hygromycin B (Invitrogen) and 3 µg/ml blasticidin S HCl. Human neuroblastoma IMR-32 cells were a gift from Igor Stagljar (CCBR, Toronto) and were maintained in MEM Eagle medium (Sigma), supplemented with 10% FBS, 1% non-essential amino acids (NEAA, Gibco) and 1% sodium pyruvate (Gibco). Cell lines were maintained at 37°C in a humidified atmosphere at 5% CO_2_. All cell lines were monitored for mycoplasma contamination using the MycoAlert mycoplasma detection kit (Lonza). Transfection of HEK 293T cells was performed using the calcium phosphate method. Media was replaced 24 h later and at 44 h post-transfection, cell lysates were collected and subjected to immunoblot analysis or co-immunoprecipitation using antibodies listed in Table S3.

### Primary murine cortical cell cultures, transfection and measurement of neurite lengths

Primary cortical neurons were obtained from E16-16.5 mouse embryos as previously described (Xu, Cobos et al., 2004). Briefly, embryonic brains were collected in Hank’s balanced salt solution (HBSS, Gibco). Cortices were dissected into small pieces, digested in 0.25% trypsin (Sigma) and the reaction was stopped by addition of DMEM supplemented with 10% FBS. Tissues were then mechanically dissociated into a single cell suspension with a glass pipette, which was then filtered through cell strainer. Dissociated primary cortical neurons were plated on tissue culture plates or chambers, coated with poly-L-lysine (20 µg/ml, Sigma) and laminin (2 µg/ml). Neurons were cultured in serum-free Neurobasal medium (Gibco) supplemented with 2% B27 (Invitrogen), 1% N2, L-glutamine, 50 IU/ml penicillin and 50 µg/ml streptomycin.

Transfection of dissociated neurons was performed using the AMAXA Nucleofector system (program O.005) with the mouse neuron Nucleofector kit (Lonza) following the manufacturer’s specifications. Briefly, 3-5 x 10^6^ cells were transfected with 2 µg GFP expressing plasmid (EGFP) as a reporter and 4 µg of plasmid DNA or 50 nM of siRNAs. All DNAs were prepared using Endo Free Maxiprep kit (Qiagen). Media was replaced 3 to 4 hours after plating. For quantification, neurons were stained for Tau-1 (axonal marker) and MAP2 (dendritic marker) and neurites, positive for Tau-1 and negative for MAP2 were considered axons. In experiments where an axonal marker was not used (ie Tuj1, phalloidin, GFP), an axon was defined as the longest neurite that was 3.5 times longer that the diameter of the soma. All measurements were performed using Volocity software (Perkin Elmer). Data are expressed as percentage of all neurons for a minimum 100 neurons per condition from 3 independent experiments unless otherwise indicated.

### Small molecule inhibitors and ligand stimulation

Cortical neurons were treated with inhibitors or ligands 4 h after plating and were fixed 33-38 h after treatment for analysis. For IGF-1R, inhibitors or activators were added in starvation medium (serum-free Neurobasal medium, supplemented with 2% B27 minus insulin, 1% L-glutamine, 50 IU/ml penicillin and 50 µg/ml streptomycin). The following inhibitors and ligands were used: TAE684 (NVP-TAE684, Selleckchem, #1108), crizotinib (PR-02341066, Selleckchem #1068), LY 294002 (Tocris), BKM120 (Buparlisib) (Selleckchem, #S2247), AG1024 (Selleckchem #S1234), PPP (CAS 477-47-4, Santa Cruz, scbt #24008) and recombinant murine IGF-1 (Peprotech).

ALKAL2-containing conditioned medium (ACM) was generated as described previously (Guan et al., 2015) with slight modifications. Briefly, HEK293T cells, in 10 cm plates, were transfected with 8.0 µg pcDNA3-FAM150B-HA using the calcium phosphate method. After 16 h, media was replaced with starvation medium (DMEM, supplemented with 0.2% FBS, 50 IU/ml penicillin and 50 µg/ml streptomycin) and ALKAL2-conditioned medium was collected after 36 h. Cortical neurons were treated with CCM and ACM, supplemented with 2% B27 and 1% N2. Activation of wild-type endogenous ALK was confirmed in IMR-32 human neuroblastoma cells.

### Protein extraction, immunoblotting, immunoprecipitation and binding assays

Cultured cells and primary cortical neurons were lysed on ice in lysis buffer (50 mM Tris, pH 7.4, 150 mM NaCl, 1 mM EDTA and 1% Triton X-100), supplemented with phosphatase and protease inhibitors(Bao, Christova et al., 2012). Embryonic brains from wild-type or knockout mice were homogenized on ice in lysis buffer and cleared supernatants were used for immunoblot analysis. For immunoblotting, protein concentrations were determined using the Bradford assay (BioRad) prior to SDS-PAGE. Antibodies used are listed in Supplementary Table 3. For co-immunoprecipitation studies, lysates were incubated with the primary antibody at 4°C for 2 h or overnight and then incubated with protein G- or protein A-Sepharose (GE Healthcare) for 2 h. Samples were washed with wash buffer (50 mM Tris, pH 7.4, 150 mM NaCl, 1 mM EDTA and 0.1% Triton X-100) and were resuspended in SDS sample buffer. For binding assays, recombinant GST (SignalChem #G52-30U) and GST-LTK (SignalChem #L11-11G) or GST-ALK (SignalChem #A19-116) were incubated with GSH-Sepharose beads (GE Healthcare) in 400 µl binding buffer (50 mM Tris, pH 7.4, 150 mM NaCl, 1 mM EDTA and 0.5% NP-40) for 1 h at 4°C. Immobilized recombinant proteins were then incubated with 400 µl embryonic brain homogenates or with recombinant His-IGF-1R (SignalChem #I02-11H) for 2 h. Samples were washed 3 times with binding buffer and were resuspended in SDS sample buffer.

### *In vitro* kinase assay

Recombinant GST-tagged LTK (SignalChem #L11-11G) was incubated in the absence or in the presence of an equal amount of recombinant His-tagged IGF-1R (SignalChem #I02-11H) in kinase buffer (50 mM HEPES, pH 7.5, 100 mM NaCl, 2 mM DTT, 5 mM MgCl_2_), supplemented with 150 µM cold ATP and 10 µCi [γ-^32^P] ATP (Perkin Elmer) for 30 min at 30°C. Reactions were terminated with SDS loading buffer. HEK293T cells were transfected with Flag-tagged IGF-1R alone or together with HA-tagged LTK. Flag immunoprecipitates of IGF-1R-Flag were incubated with Protein G Sepharose beads (GE Healthcare). Beads were pelleted, washed 3 times in wash buffer and 3 times in kinase buffer. Kinase reactions were performed for 30 min at 30°C in kinase buffer, supplemented with 150 µM cold ATP and with 10 µCi [γ-^32^P] ATP (Perkin Elmer). Reactions were stopped by addition of gel loading buffer and were separated by SDS-PAGE. Gels were dried and the phosphorylation products of kinase reactions were detected by phosphorimaging.

### Cell surface biotinylation assay

High density cortical neuron cultures or T-REx-LTK cells were cooled on ice to prevent receptor endocytosis and were rinsed with PBS supplemented with 0.5 mM MgCl_2_ and 1 mM CaCl_2_. Cell-surface expression level of IGF-1R was examined using membrane-impermeable biotin EZ-Link Sulfo-NHS-LC-biotin (Pierce). Cells were incubated with 1 mg/ml biotin for 20 min on ice and unreacted biotin was removed by three washes with a quenching agent (50 mM glycine and 0.5% BSA in PBS with MgCl_2_ and CaCl_2_). Cells were lysed in RIPA buffer containing protease and phosphatase inhibitors and biotinylated proteins were precipitated with NeutrAvidin agarose resin (Pierce). In some experiments, cells were incubated with CCM or ACM on ice or at 37°C to allow endocytosis and then were lysed and precipitated with NeutrAvidin agarose beads. Agarose beads were washed twice in RIPA containing 500 mM NaCl and once in standard RIPA buffer. Biotinylated surface proteins and total lysates were subjected to immunoblotting and quantitated using Image J software (NIH, Bethesda).

### RNA extraction and Real-time PCR

Total RNA was extracted from primary cortical neurons or brain tissues using Pure Link RNA Mini Kit (Life Technologies) and reverse transcribed into cDNA using oligo dT primers and M-MLV Reverse Transcriptase (Invitrogen). Real-time PCR was performed on Quant Studio 6 Flex Real-time PCR system (Applied Biosystems) using SYBR Green Master mix (Applied Biosystems). Relative gene expression was quantified by ΔΔCt method and normalized to HPRT1 or GAPDH. The validated primer sequences are listed in Table S2.

### Immunocytochemistry

Neuronal cultures were fixed using 4% (w/v) PFA for 15 min except for cultures used for Tau-1 staining where fixation was performed for 30 min. Fixed cells were permeabilized with 0.2% Triton X-100 in PBS for 5 min, washed with PBS and blocked at room temperature for 1 h using either 3% bovine serum albumin (BSA, Sigma) or 5% goat serum according to the primary antibody used. Cells were incubated overnight at 4°C with primary antibodies, washed twice with PBS and incubated with secondary antibodies at room temperature for 1 h. Cells were washed and directly visualized or were fixed and mounted with ProLong Gold (ThermoFisher). Images were taken in a random fashion using a Zeiss inverted microscope equipped with CCD camera (Hamamatsu Photonic Systems) and Volocity software (Perkin Elmer).

### Histology and staining of brain sections

Mice were anesthetized with 2.5% Avertin (2,2,2-tribromethanol, Sigma) and perfused transcardially with 4% paraformaldehyde in 0.1M Na-phosphate buffer, pH7.4 (PFA). Brains were removed and stored at 4°C overnight in fixative, and then for 1 day at 4°C in 30% sucrose in 0.1M phosphate buffer, pH 7.4. Brains were cryosectioned using Leica CM3050S cryostat and processed for staining. For Nissl staining, cryosectioned brain slices on slides were immersed in 1:1 alcohol/chloroform at room temperature overnight and then were rehydrated sequentially through 100%, 95% alcohol and then distilled water. Slides were stained with 0.1% cresyl violet solution for 3-5 minutes, rinsed quickly in distilled water, developed in 95% ethyl alcohol up to 20 min, dehydrated in 100% alcohol twice for 5 min each, cleared using xylene twice for 5 min each and then mounted with permanent mounting medium. For immunohistochemistry, brain sections were stained as previously described (Polleux & Ghosh, 2002) using antibodies listed in Table S3. For quantitation in P2 and P7 mice, cells located within each of 8 equally sized bins spanning the cortex were counted and were plotted as a percent of DAPI+ cells. For Golgi staining, brains, from adult mice, at least 3 per genotype, were impregnated with Golgi-Cox solution at room temperature in the dark for 30 days using the FD Rapid Golgi Stain Kit (FD Neurotechnologies). Brains were then transferred to solution C for 2 days, followed by cryosectioning at 60 µm and staining according to the manufacturer’s protocol. Pyramidal neurons in layer V of the neocortex and the callosal tract were captured by brightfield microscopy.

### *In vivo* migration assays using BrdU labelling

E14.5 pregnant wild-type, *Alk^-/-^, Ltk^-/-^* and *DKO* mice were labeled *in vivo* by intraperitoneal injections with BrdU (150 µg/g body weight, Sigma-Aldrich,). In some experiments PPP (20 mg/kg body weight) (Girnita, Girnita et al., 2004) or saline (as control) were injected 24 h after BrdU. E17.5 embryos, P2 and P7 pups were sacrificed and brains were fixed in 4% PFA and cryoprotected in 30% (w/v) sucrose at 4°C overnight. Serial cryosections (12 µm) were incubated with anti-BrdU antibody at 4°C overnight. For quantitation in P2 and P7 mice, BrdU+ cells located within each of 8 equally sized bins spanning the cortex were counted.

### *In Situ* Hybridization

Single-molecule fluorescence *in situ* hybridization (smFISH) was performed using Multiplex Fluorescent Reagent Kit (v2) according to manufacturer’s instructions (Advanced Cell Diagnostics), with probes targeting *Alk* (cat# 501131-C1, *Ltk* (cat# 530801-C2) and *Alkal2* (cat# 531801-C3). Positive (POLR2A: Channel C1, PPIB: Channel C2, UBC: Channel C3: cat#320881) and negative (Bacterial dap gene: cat# 320871) probes were used to validate specificity. Briefly, embryonic (E15-16) or transcardially perfused adult brains were fixed in 4% PFA for 24 h, cryoprotected in 30% (w/v) sucrose for 24 h at 4°C, and coronally cryosectioned at 20 µm. Sections were treated with target retrieval reagent for 5 min, followed by 40°C in proteases plus permeabilization for 10 min for embryonic brain and 30 min for adult brain. Sections were immunostained using antibodies listed in Table S3. Confocal images were acquired using a Nikon Ti2 inverted confocal microscope and analyzed using Volocity software (Perkin Elmer). Widefield images were acquired using Zeiss Axio Scan Z1 and analyzed with Zeiss Zen Black software.

### Behavioral studies

Knockout and wild-type animals were tested on a battery of physical and cognitive behavioral studies. <u>Dark and Light box.</u> This assay for anxiety-related behavior and was carried out as described previously (Holmes, Yang et al., 2002). The apparatus consisted of a clear plexiglass cage (44 x 21 x 21 cm), illuminated by a 60W desk lamp overhead and a compartment made from black plexiglass with a covered top. Compartments were separated by a partition, with a small opening (12 x 5 cm) at floor level. Mice were placed in the light compartment, facing away from the partition and allowed to explore freely for 3 min. The latency to enter the dark box and the number of light-dark transitions (re-enters) and length of time in light or dark were recorded. <u>Y-maze test.</u> Spatial memory was tested in a Y-maze (Kraeuter et al., 2019, Miedel et al., 2017), consisting of three identical arms with dimensions 38 x 76 x 12.7 cm that met at the center separated by an angle of 120° between each pair of arms (San Diego Instruments). Mice were placed at the end of one arm facing the center and explorations were recorded for 8 minutes by a camera mounted above the maze and by video-tracking software (Bioserve Viewer2). Entries of all four paws, 5 cm into the arm were recorded. Alternation behavior comprising consecutive entries into each of the three arms without repetition was monitored with the percentage of spontaneous alternations (actual alternation divided by the possible alternations (ie total entries-2) x 100) were quantitated (Miedel et al., 2017). <u>Puzzle box paradigm.</u> The Puzzle box assesses cognitive flexibility and problem solving ability in which mice must leave a brightly lit, open arena to reach a dark, enclosed area through increasing complexity of barriers as previously described (Ben Abdallah et al., 2011). Each Day, three trials were performed with a 2-minute inter-trial interval for a maximum of 5 minutes per trial. On Day 1, trial 1, both the doorway and underpass were open while in trials 2 and 3, only the underpass was open. On Day 2, trial 4, only the underpass is open while in trials 5 and 6 the underpass was blocked with bedding. On Day 3, trial 7, the underpass was blocked with bedding while in trials 8 and 9, the underpass was blocked with a removable cardboard plug.

### Statistical Analysis

Statistical evaluation between two groups was performed using unpaired Student’s t tests and between more than 2 groups was carried out using one-way ANOVA with Dunnett’s post hoc analysis as indicated in the figure legends. The quantitative results are expressed as the mean +/- the standard error of the mean (SEM). For all analyses performed, significance was defined as ****p<0.0001, ***p<0.001, **p<0.01, *p< 0.05 and ns, not significant. Data analysis was done using Prism v8 (Graph Pad Software).

## Acknowledgements

We thank Dr. Stephen Morris (St. Jude CRH, Memphis) and Dr. Joseph Weiss (OHSU, Oregon) for kindly providing Alk and Ltk knockout mice, Dr. Ruth Palmer (University of Gotenburg, Sweden) for providing plasmids encoding ALKAL1 and ALKAL2 and Dr. Igor Stagljar (University of Toronto) for IMR-32 cells and Dr. Amy J. Ramsey (University of Toronto) for access to equipment for behavioural studies. The authors wish to thank Xin Zhao, Solmaz Alizadeh and Peter Poliszczuk for technical support during the early stages of the project. This work was funding by a CIHR Foundation grant (#FDN148455) to L.A.

---

**Figure S1.**
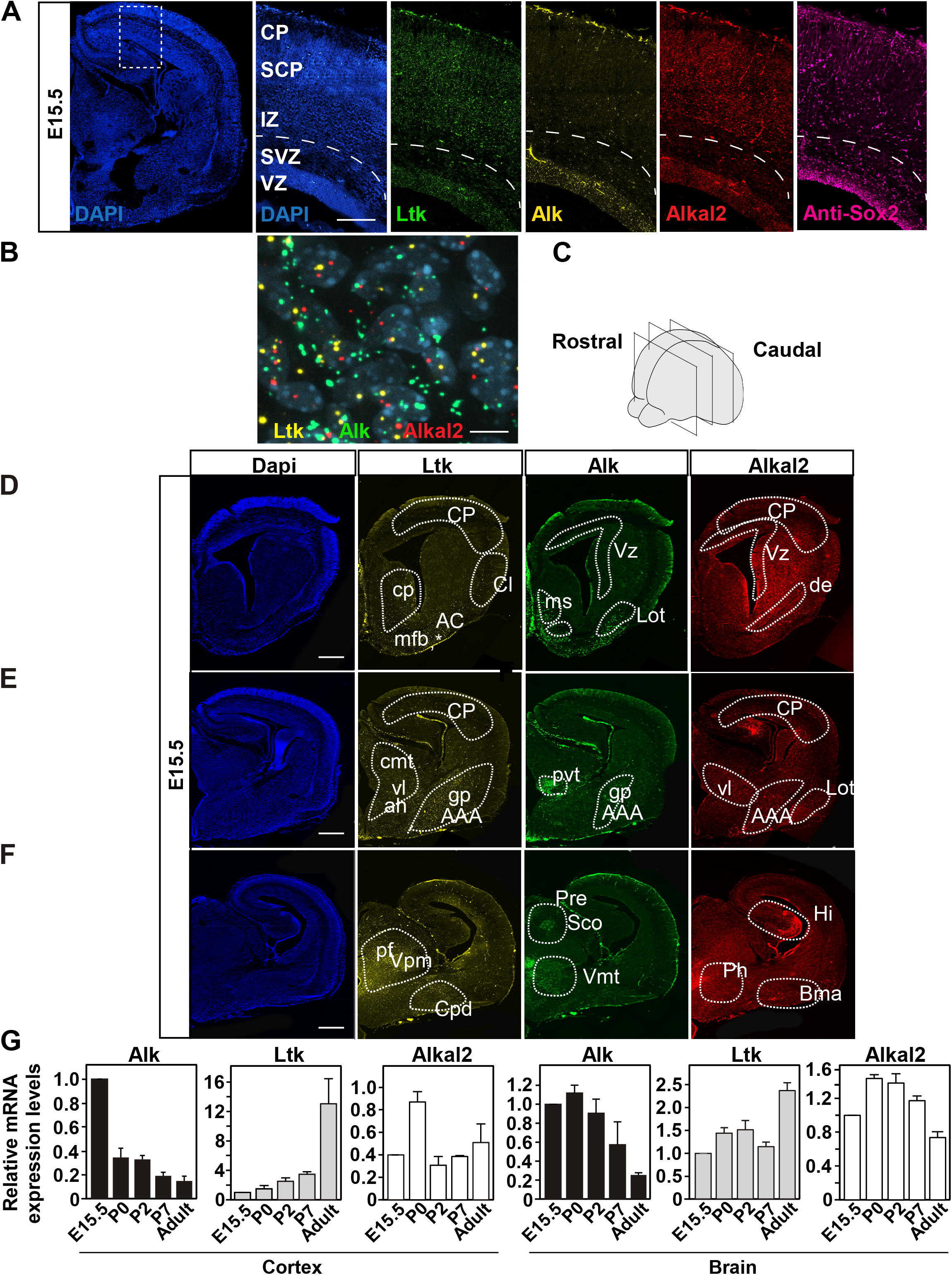
Expression of Ltk, Alk and Alkal2 in murine embryonic and adult cortex and brain. **(A)** Expression of Alk, Ltk and Alkal2 at E15.5 was determined by single molecule *in situ* analysis. A representative coronal wide-field DAPI stained image (blue) of a coronal section of an E15.5 brain (left). Higher magnifications of the inset region (right panels) show mRNA expression of Ltk (green), Alk (yellow) and the ligand, Alkal2 (red) and are counterstained with Sox2 protein (anti-Sox2: purple) and DAPI (blue). **(B)** A high magnification (63X) image from the SVZ shows mRNA expression of Alk (green), Ltk (yellow) and Alkal2 (red) counter-stained with DAPI (blue). **(C-F)** Ltk and Alk are expressed in distinct patterns in E15.5 embryos during the onset of neurogenesis and cortical plate formation. **(C)** Schematic denoting the approximate location of coronal sections E15.5 at which the expression was examined in D, E, F, rostral to caudal, respectively. **(D-F)** Ltk (yellow) is found rostrally in the neo-cortical plate (CP), claustrum (Cl), accumbens nucleus (AC), amygdaloid area (AAA) and caudate putamen (cp). High expression of Ltk was observed in thalamic area, ventral thalamic nucleus (Vl), paraventricular thalamic nucleus (Pvt), hypothalamic area (Ah) and globus pallidus (Gp). Alk (green) was prominently expressed near ventricular regions throughout the neocerebrum with robust expression in the area of the medial spectrum nucleus (Ms), lateral olfactory bulb (Lot), paraventricular thalamic nucleus (Pvt), ventromedial thalamic nucleus (Vmt), pretactal area (Pre) and subcommissural organ (Sco). Alka2 (red), is expressed in many regions, overlapping with Ltk and Alk. Co-expression was observed throughout cortical plate (CP), ventricular zone, ventrolateral thalamic nucleus (vl), lateral olfactory bulb, amygdaloid area, and hypothalamic area. **(G)** The expression levels of Alk, Ltk and Alkal2 mRNA in the embryonic, post-natal (P2 and P7) and adult (3-6 month) cortex and brain was determined by qPCR. Data is plotted as the mean +/- SEM, from 3 independent experiments.

**Figure S2.**
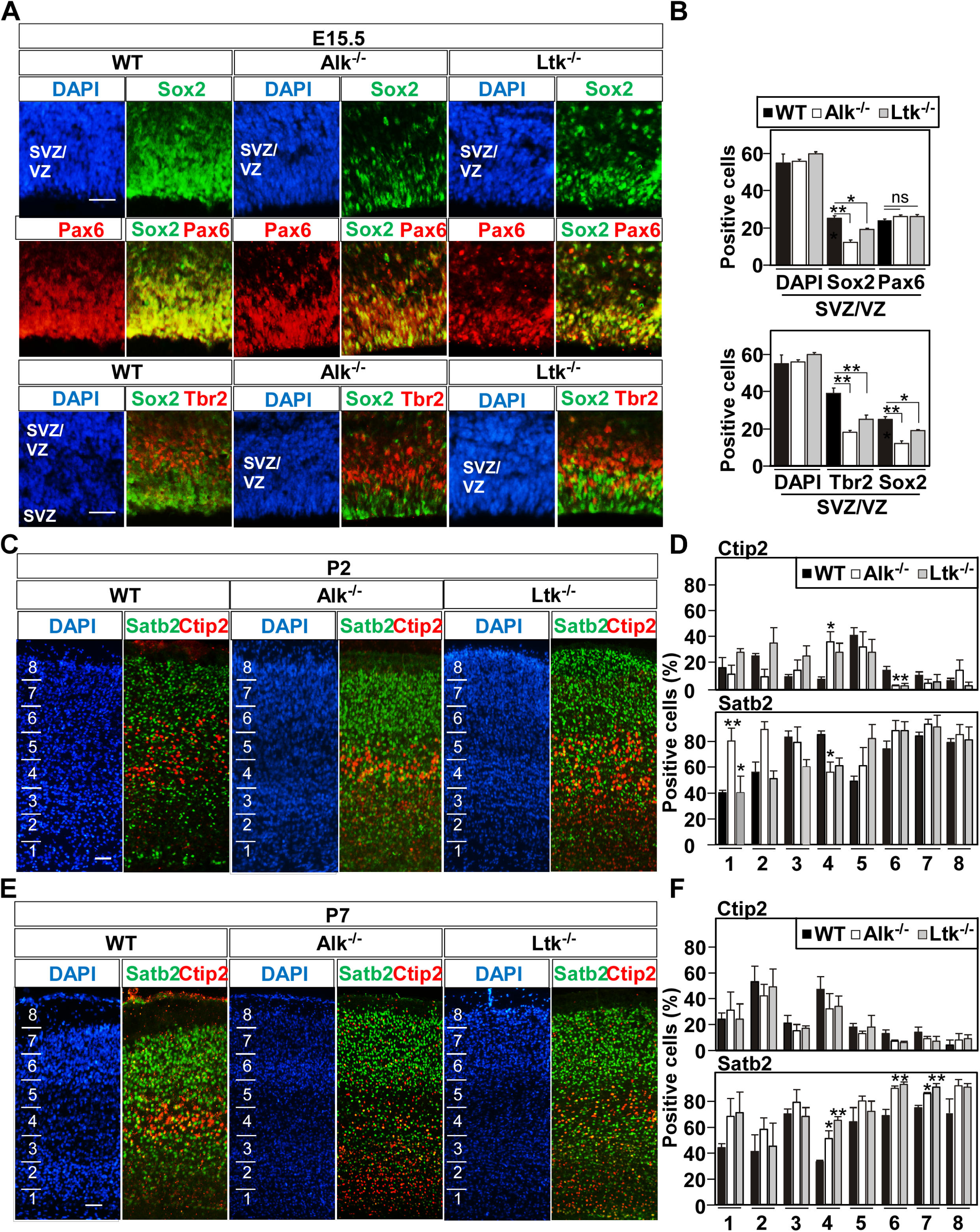
Disruption of Ltk and Alk alters early cortical plate patterning. **(A and B)** Coronal sections of E15.5 WT, Ltk^-/-^ and Alk^-/-^ developing cortices were co-stained with either anti-Sox2 (green) and anti-Pax6 (red) or with anti-Sox2 (green) and anti-Tbr2 (red) and both counter-stained with DAPI (blue). Quantitation of total DAPI+ cells and Sox2+, Pax6+ or Tbr2+ progenitor cells in the SVZ/VZ are plotted as the mean +/- SEM of 3 independent experiments. **(C-E)** Coronal sections of WT, Ltk^-/-^ and Alk^-/-^ developing cortices stained with DAPI (blue) and co-stained with anti-Ctip2 (red) and Satb2 (green) at P2 and P7. **(C and E)** Quantitation of Ctip2+ and Satb2+ progenitor cells as a percent of total DAPI+ cells in each of 8 bins (marked on left) within the cortex (dashed lines) are plotted as the mean +/- SEM of 3 independent experiments. Statistical significance: **p<0.01, *p<0.05 by one-way ANOVA using Dunnett’s test. The location of the cortical plate (CP), intermediate zone (IZ), subventricular/ventricular zone SVZ/VZ in P2 mice and Layers I-VI in P7 WT mice are indicated. Scale bar, 50 µm.

**Figure S3.**
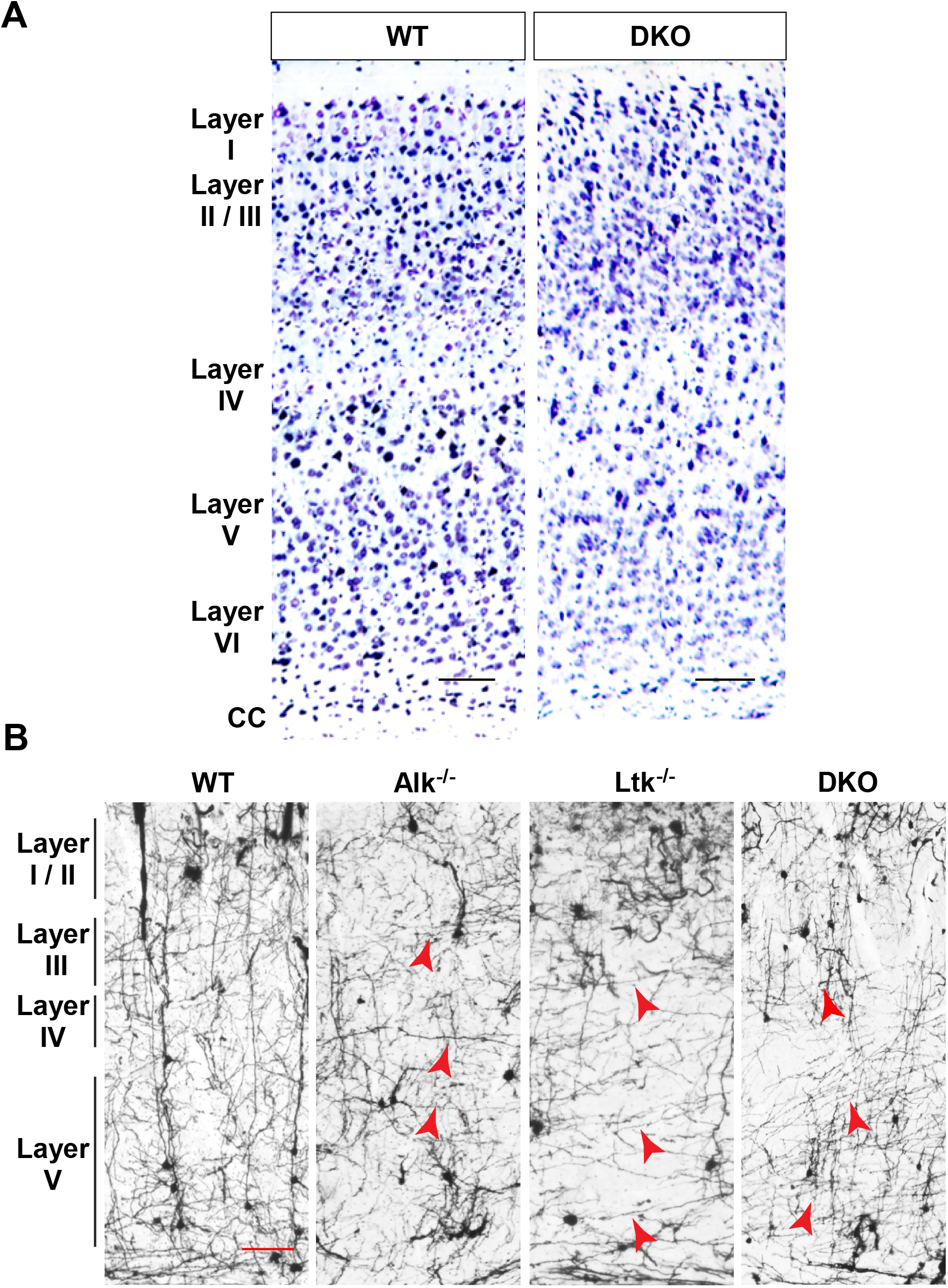
Analysis of Coronal sections of WT and Ltk/Alk knockout mice by Nissl and Golgi staining. **(A)** Analysis of cortical layering in vivo by Nissl staining of adult WT and DKO murine brains. Nissl-stained coronal sections reveal a similar pattern of cortical layering in wild type (WT) and DKO adults. Scale Bar = 100 µm. **(B)** Golgi stains of adult cortices in wild-type (WT), single (Ltk-/-and Alk-/-) and double (DKO) knockout mice show defects in neuronal projections. Representative coronal sections obtained from an independent experiment from those in Fig. 3E are shown. Examples of aberrant projections in pyramidal neurons in knockout mice are marked (red arrowheads).

**Figure S4.**
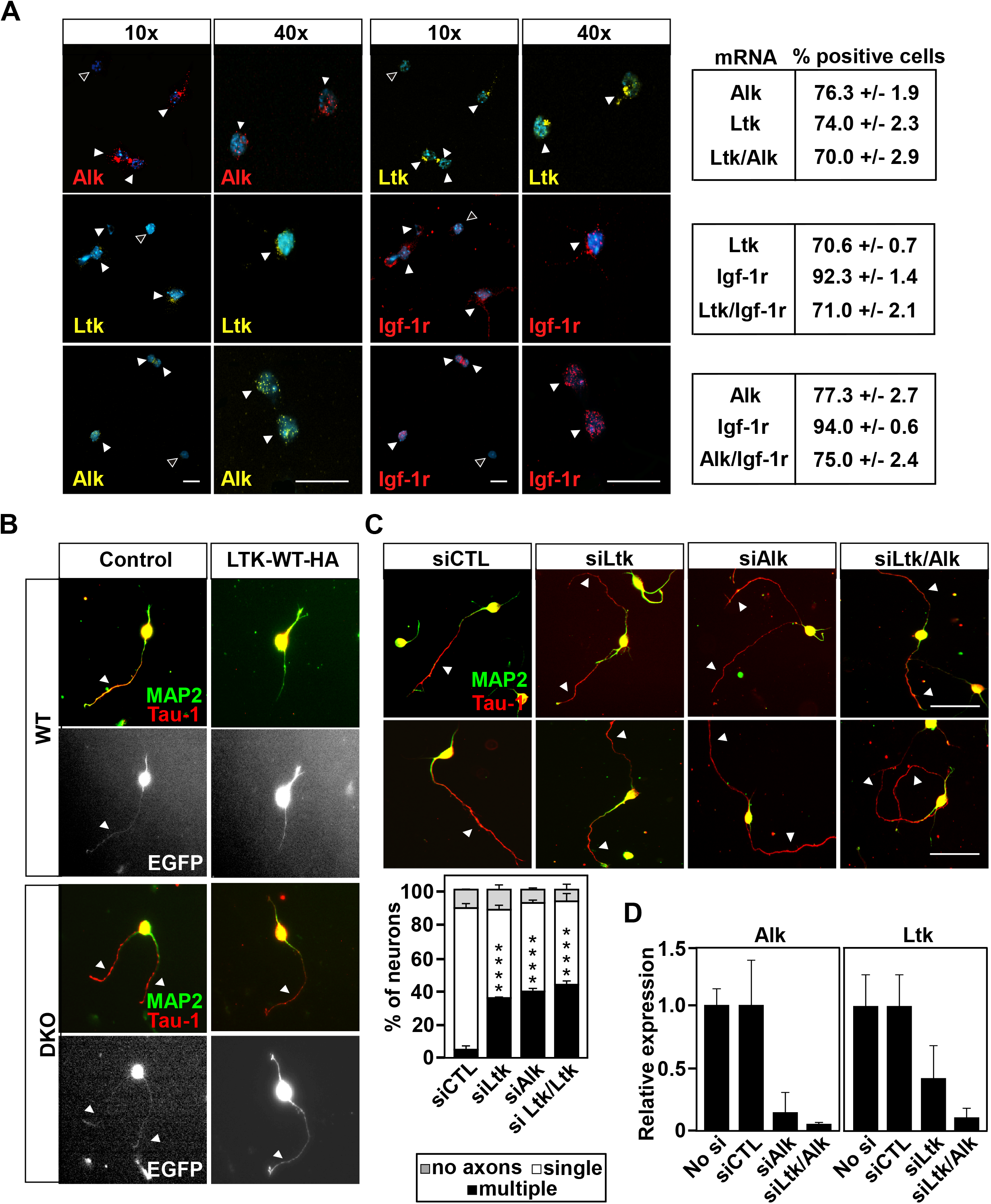
Change of expression or activity of Ltk and Alk results in defects in neuronal polarity. **(A)** Analysis of mRNA expression of Ltk, Alk and Igf-1r in isolated neurons. Expression of Alk, Ltk and Igf-1r in isolated primary cortical neurons from WT mice was determined by single molecule *in situ* analysis. Independent images taken at 10X and 40X magnification are shown. Filled arrowheads mark positive cells and open arrowheads mark negative cells. Quantitation of the percent of positive cells from 3 experiments and a minimum of 50 cells per condition was tabulated (right). **(B)** Additional representative images of dissociated WT and DKO cortical neurons, electroporated with plasmids encoding EGFP alone (control) or together with HA-tagged LTK-WT (as in Figure 3C) are shown. Cells were fixed 38 h after plating and then stained for Tau-1 (axons, red)) and MAP2 (dendrites, green) with transfected cells identified by EGFP staining (white). **(C)** Abrogation of the expression of Ltk, Alk or both in cortical neurons using siRNAs results in defects similar to that observed in neurons isolated from knockout mice. WT cortical neurons were transfected with siCtl, siLtk and/or siAlk. Representative images of cells fixed at 38h and stained with Tau-1 (axons, red) and MAP2 (dendrites, green) are shown. Quantitation of the percent of neurons with multiple, single or no axons is plotted as the mean ± SEM of n=150 neurons from 3 independent experiments. Scale bar = 20 µm. Statistical significance: ****p<0.0001 by one-way ANOVA using Dunnett’s test. **(D)** Knockdown efficiency in neurons was evaluated by real-time PCR. Data are plotted as mean ± SD from 3 independent experiments.

**Figure S5.**
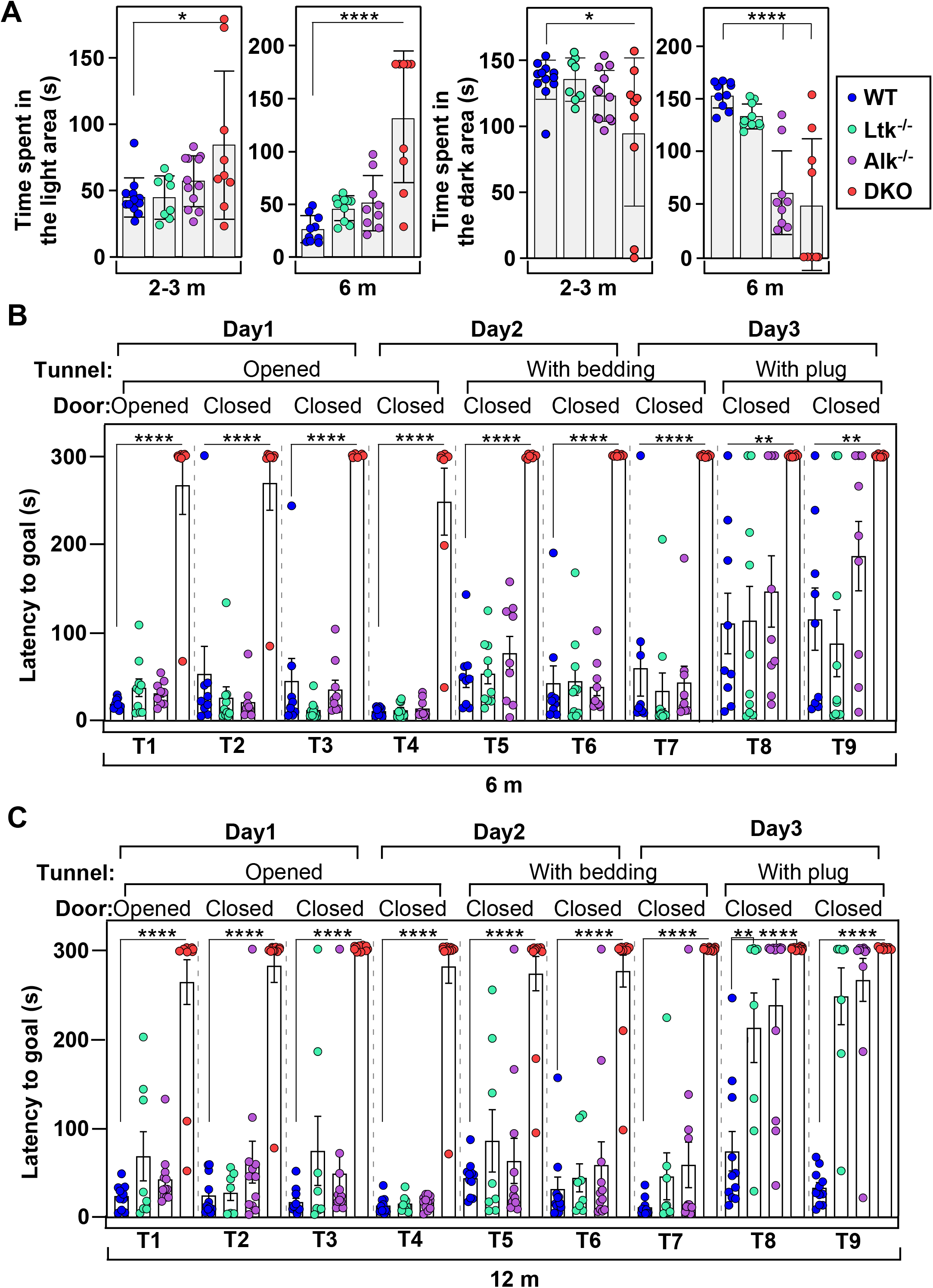
Performance of mice in light-dark box and in puzzle box tests. **(A)** Light-dark box. Time spent in the light or dark areas of individual mice of 2-3 or 6 months of age of the indicated genotypes is plotted. **(B)** Puzzle-box test. Latency to reach the goal box during the 9 trials of the test is plotted for each individual 6 (B) or 12 **(C)** month old mouse of the indicated genotypes. Results in all panels are plotted as mean ± SEM. Data were analyzed using one-way ANOVA and comparisons to WT were performed using Dunnett’s test (****p<0.0001, ***p<0.001, **p<0.01, *p<0.05, ns, not significant).

**Figure S6.**
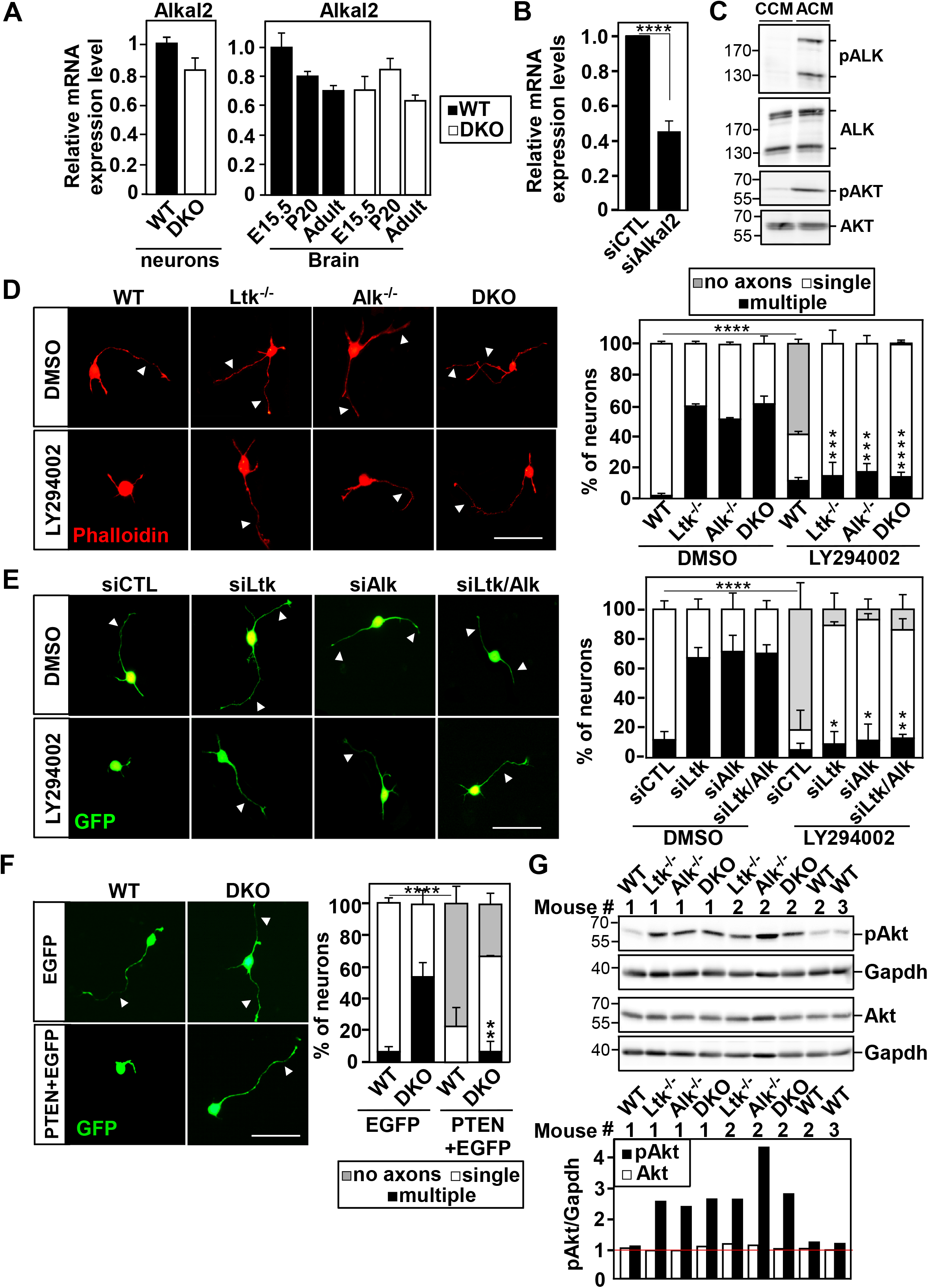
ALKAL2 ligand-mediated activation of LTK/ALK results in defects in neuronal polarity. PI3 kinase is activated in cortical neurons from knockout mice. (**A**) The expression of Alkal2 mRNA in E16 dissociated cortical neurons or in embryonic, P20 and adult (3-6 month) brain in WT and DKO mice was determined by qPCR. Data is plotted as the mean +/- SD, from 3 independent experiments. (**B**) Alkal2 knockdown efficiency was determined by real-time PCR. Data are plotted as mean ± SD of 3 independent experiments. (**C**) IMR-32 cells were treated for 20 min with control (CCM) or ALKAL2 (ACM) conditioned media. Representative immunoblots indicating the levels of pALK (pALK^Y1586^), pAKT (pAKT^S473^) and total ALK and AKT in cell lysates are shown. The increased levels of phosphorylated ALK and AKT indicate that ligand addition increased the activity of ALK and ALK downstream signaling. (**D**) Treatment of cortical neurons from WT or KO mice with PI3 kinase inhibitors reverses polarity defects caused by loss of Ltk/Alk. Dissociated E16 cortical neurons were treated with LY294002 (10 µM) or DMSO control, 4 h after plating and then fixed at 33 h and stained with Phalloidin. Arrowheads mark the axon-like projections. Scale bars are 20 µm. Quantitation of neurons with multiple, single or no axon-like projections is plotted as the mean ± SD, from three independent experiments. (**E**) Inhibition of PI3K reverses the effect of siRNA-mediated knockdown of Ltk/Alk in cortical neurons. Representative images of dissociated E16 WT cortical neurons co-transfected with the indicated siRNAs and GFP were treated with LY294002 (PI3K inhibitor) 6 h after plating and then fixed at 33 h. Scale bar = 20 µm. Quantitation of the percent of neurons with multiple, single or no axon-like projections is plotted as mean ± SEM of 3 independent experiments. (**F**) Ectopic expression of PTEN rescues the multiple axon phenotype in cortical neurons. Representative images of cortical neurons, isolated from WT or DKO embryos and transfected with plasmids encoding GFP alone or HA-tagged PTEN. Scale bar = 20 µm. The percentage of neurons in GFP-positive cells with multiple, single or no axon-like projections is plotted as the mean ± SEM (n=100 neurons) from 3 independent experiments. (**G**) Analysis of brain homogenates reveal activated downstream PI3 kinase signaling in DKO mice. Brain lysates from individual mice of the indicated genotypes were subjected to immunoblotting with antibodies against phosphorylated AKT (pAKT^S473^) and AKT. Blots were re-probed with GAPDH antibody as a loading control. Quantitation of band intensities for pAkt and Akt were normalized to those of Gapdh in each lane and the fold change relative to control WT1 (WT, mouse #1) is plotted. Data in **D-E** were analyzed using one-way ANOVA and comparisons to the concurrent DMSO control were performed using Dunnett’s test. Data in **B** and **F** were analysed using two-tailed Student’s t-test. Statistical significance: ****p<0.0001, ***p<0.001, **p<0.01, *p<0.05.

**Figure S7.**
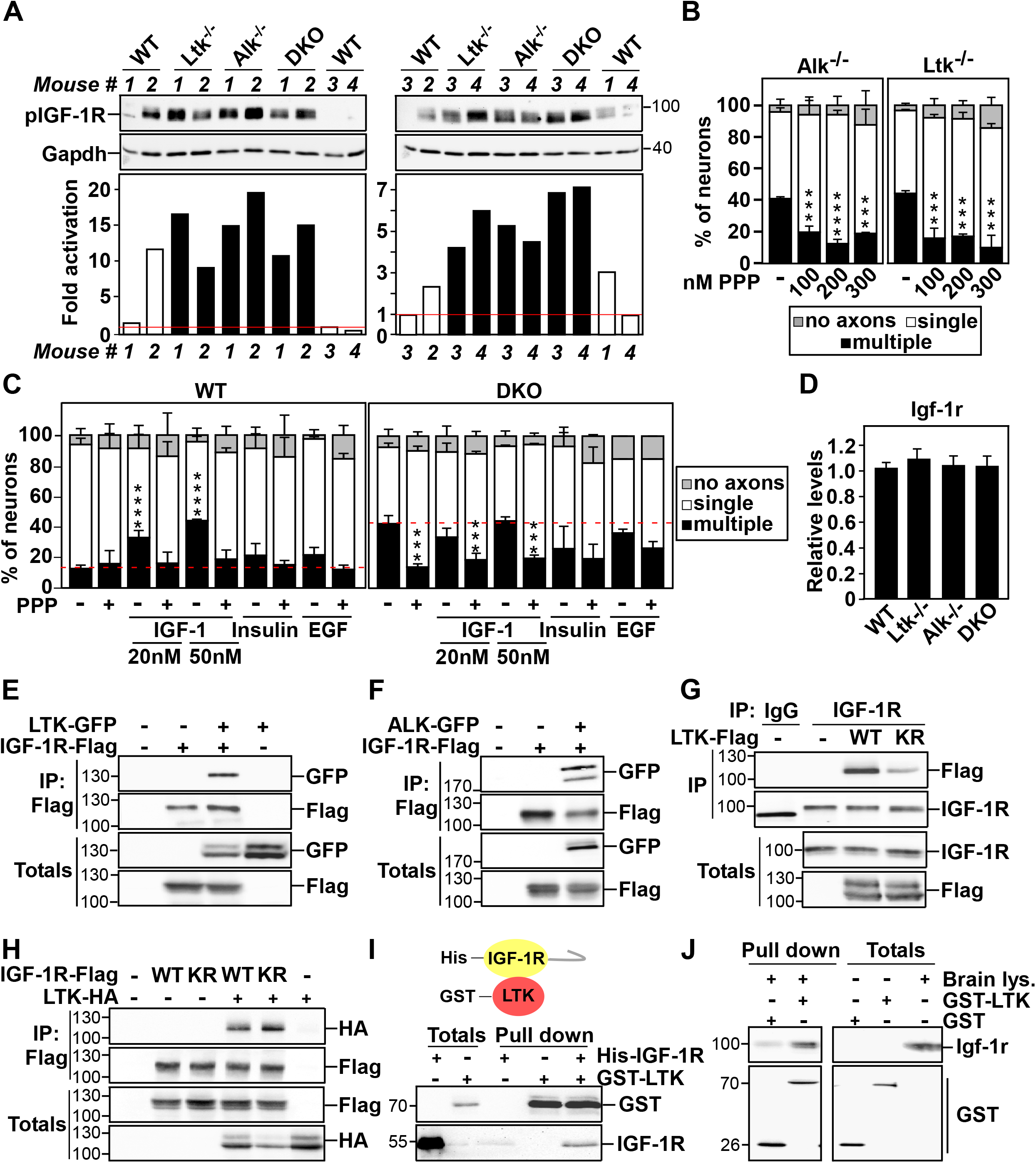
LTK/ALK interacts with IGF-1R and modulates its activity. Loss of LTK/ALK promotes IGF-1R activity. **(A)** Brain lysates from 4 individual mice of the indicated genotypes were subjected to immunoblotting with phospho-IGF-1R (pIGF-1R^Y1135/1136^) and Gapdh as control to verify equal protein loading. Quantitation of band intensities for pIGF-1R/Gapdh as fold change relative to control WT3 (WT, mouse #3) is plotted. **(B)** The IGF-1R inhibitor, PPP rescues the multiple axon phenotype in cortical neurons from Alk or Ltk knockout mice. Cortical neurons, isolated from Alk or Ltk knockout mice were treated with varying concentrations of PPP or DMSO as control 4h after plating, fixed and stained for Tuj1. Quantitation of the percent of neurons with multiple, single or no axon-like projections is plotted as the mean ± SD, n = 100 neurons from 2 independent experiments. **(C)** Analysis of the effect of ectopically applied recombinant IGF-1. Cortical neurons isolated from WT or DKO mice electroporated with a GFP plasmid were treated with PPP (200 nM), DMSO as control and /or IGF-1 (20 or 50 nM), insulin (100 nM) or EGF (20 ng/ml). Cells were fixed at 33h and GFP positive cells were analyzed. Quantitation of the percent of neurons with multiple, single or no axon-like projections is plotted as the mean ± SD, n=100 neurons from 2 (for EGF) or n=200 neurons from 4 (remainder) independent experiments. Data in **B** and **C** were analyzed using one-way ANOVA and comparisons to control were performed using Dunnett’s test (**** p<0.0001, *** p<0.001). **(D)** Expression of Igf-1r mRNA was determined by qPCR in cortical neurons from WT and knockout mice. Data are plotted as mean ± SD of 3 independent experiments. **(E-F)** IGF-1R associates with LTK and ALK in HEK293T cells. Lysates from HEK 293T cells, transfected with IGF-1R-Flag, LTK-GFP **(E)** or ALK-GFP **(F)** were subjected to immunoprecipitation using anti-Flag antibody and co-precipitated LTK **(E)** or ALK **(F)** was detected by immunoblotting using anti-GFP antibody. **(G)** LTK activity is required for interaction with IGF-1R. HEK293T cells were transfected with wild type (WT) or kinase inactive (KR) versions of LTK-Flag. Endogenous IGF-1R was immunoprecipitated with anti-IGF-1R antibody and co-precipitated LTK-Flag was detected by immunoblotting with anti-Flag antibody. **(H)** LTK/ALK interacts with WT and kinase inactive (KR) variants of IGF-1R. Lysates from HEK293T cells, co-transfected with LTK-WT-HA and WT or KR versions of IGF-1R-Flag were subjected to immunoprecipitation with anti-Flag antibody and co-precipitated LTK was detected by immunoblotting with anti-HA antibody. **(I)** GST-LTK immobilized to GSH-Sepharose beads was incubated with recombinant His-IGF-1R. Pull down and totals were subjected to immunoblotting using indicated antibodies. **(J)** Purified recombinant GST-LTK interacts with endogenous Igf-1r. Lysates from WT embryonic brains were incubated with GST-LTK or GST alone, immobilized to GSH-Sepharose beads. Bound and total proteins were subjected to immunoblotting with anti-IGF-1R antibody. **(E-J)** A representative result from 3 independent experiments is shown.

**Supplementary Table 1:**
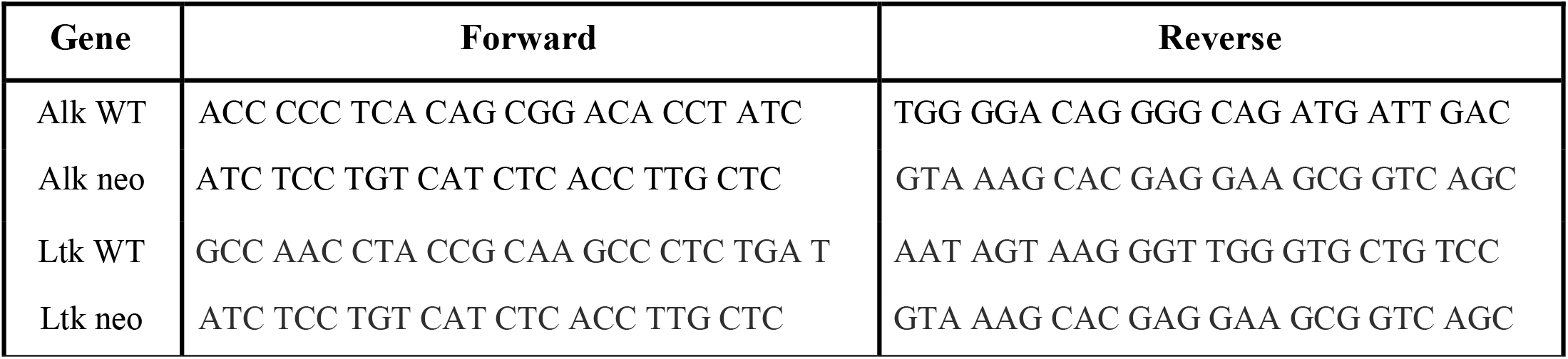
Sequences of primers used for genotyping

**Supplementary Table 2:**
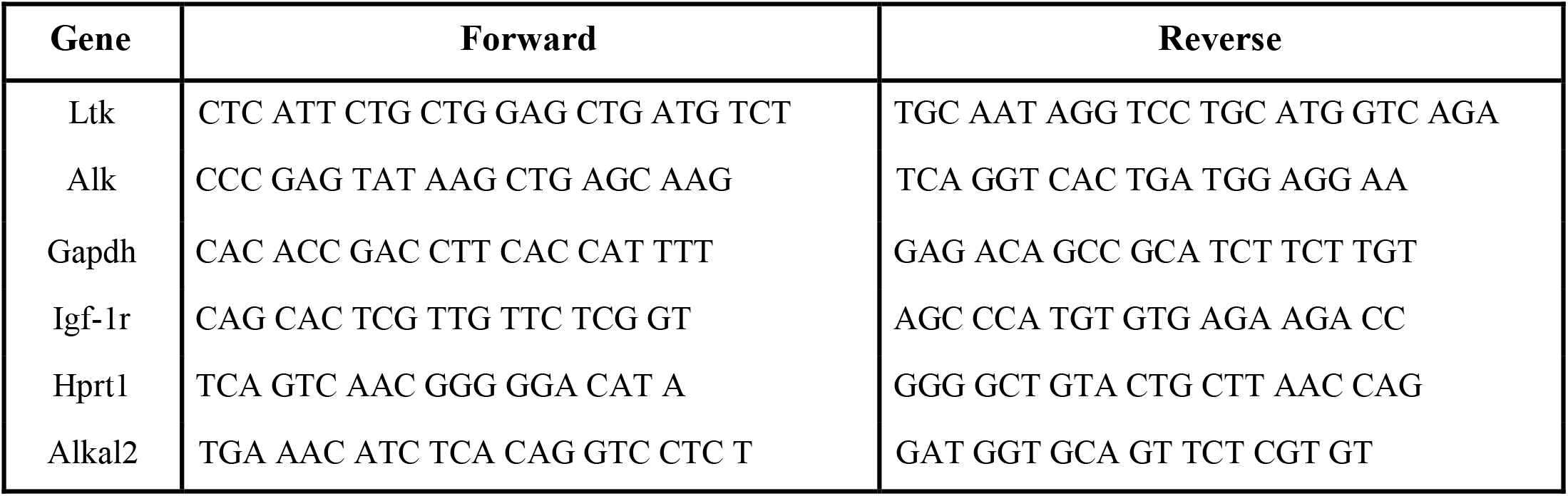
Sequences of primers used for Real-Time PCR

**Supplementary Table 3:** Antibody list

| **Antibody** | **Host** | **Application** | **Dilution** | **Suppliers** | **Catalog number** |
| --- | --- | --- | --- | --- | --- |
| MAP2 | Mouse | ICC | 1:400 | Millipore | MAB3418 |
| NeuN | Rabbit | IHC | 1:500 | Cell Signaling | 12943 |
| Tau-1 (PC1C6) | Mouse | ICC | 1:200 | Millipore | MAB3420 |
| SOX2 | Mouse | IHC | 1:50 | Santa Cruz | sc-365823 |
| TBR2 | Rabbit | IHC | 1:200 | Abcam | ab23345 |
| SATB2 | Rabbit | IHC | 1:500 | Abcam | ab51502 |
| PAX6 | Rabbit | IHC | 1:100 | Covance | PRB-278P |
| TUJ-1 | Mouse | ICC | 1:500 | BioLegend | MMS-435P |
| BrDU | Rat | IHC | 1:1000 | Abcam | ab142567 |
| CTIP2 | Rat | IHC | 1:300 | Abcam | ab18465 |
| ALK (D5F3) | Rabbit | WB | 1:1000 | Cell Signaling | 3633 |
| ALK (D5F3) | Rabbit | IP | 1:100 | Cell Signaling | 3633 |
| pALK (Tyr 1586) | Rabbit | WB | 1:1000 | Cell Signaling | 3348 |
| pAKT (Ser 473) | Rabbit | WB | 1:1000 | Cell Signaling | 9271 |
| AKT | Rabbit | WB | 1:1000 | Cell Signaling | 9272 |
| GSK3β (27C10) | Rabbit | WB | 1:1000 | Cell Signaling | 9315 |
| pGSK3β (Ser 9) | Rabbit | WB | 1:1000 | Cell Signaling | 9323 |
| IGF-1Rβ (D23H3) | Rabbit | WB | 1:1000 | Cell Signaling | 9750 |
| IGF-1Rβ (D23H3) | Rabbit | IP | 1:100 | Cell Signaling | 9750 |
| pIGF-1Rβ (Tyr 1135/1136) | Rabbit | WB | 1:1000 | Cell Signaling | 3024 |
| IGF-1Rα (24-60) | Mouse | NA | 1.5 μg/μl | ThermoFisher | MA5-13817 |
| Flag M2 | Mouse | WB | 1:3000 | Sigma | F3165 |
| Flag M2 | Mouse | IP | 1:1000 | Sigma | F3165 |
| HA | Rat | WB | 1:1000 | Roche | 11867423001 |
| GST | Rabbit | WB | 1:1000 | Cell Signaling | 2625 |
| GFP | Rabbit | WB | 1:1000 | Cell Signaling | 2555 |
| GFP | Mouse | IP | 1:100 | Santa Cruz | sc9996 |
| GAPDH | Rabbit | WB | 1:10000 | Sigma | G9545 |

